# A high-resolution dimeric structure reveals a critical role of α-helix 9 in apoptotic Bax pore assembly in the mitochondrial membrane

**DOI:** 10.64898/2025.12.30.697096

**Authors:** Zhi Zhang, Liujuan Zhou, Chenyi Liao, Fujiao Lv, Fei Qi, Lingyu Du, Justin Pogmore, Juan del Rosaio, David W. Andrews, Bo OuYang, Jialing Lin

**Author notes:** These authors contributed equally to this work as first authors (Zhi Zhang, Liujuan Zhou, Chenyi Liao). Correspondence: Jialing Lin, Department of Biochemistry and Physiology, University of Oklahoma Health Campus, 940 Stanton L. Young Boulevard, BMSB 935, Oklahoma City, Oklahoma 73104, United States of America; Bo OuYang, Shanghai Institute of Biochemistry and Cell Biology, CAS Center for Excellence in Molecular Cell Science, Chinese Academy of Sciences, 333 Haike Road, Shanghai, China 201203.

## Abstract

Bax functions as a proapoptotic protein perforating the mitochondrial membrane to release mitochondrial proteins and DNA that kill the cell. Structures of full-length Bax monomer in solution and core domain dimers in solution and bound to lipid/detergent bicelles reveal how Bax proteins must be activated, unfold, refold and dimerize prior to forming oligomeric pores in the membrane. In particular, amphipathic core dimer forms part of the pore wall between the nonpolar lipid bilayer and the aqueous conduit. However, atomic resolution structures of other parts of the pore were scarce. Here, we elucidate a structure of Bax C-terminal α-helix 9 (α9) in lysolipid micelles using NMR. According to this high-resolution structure the α9 regions form an amphipathic helical dimer with an extended nonpolar interface and several uncharged polar residues on the surface. Structure-guided mutagenesis and functional assessment demonstrate that the nonpolar interactions are important for Bax dimerization in and perforation of the mitochondrial membrane. Surprisingly the polar residues are also important because they form bifurcated hydrogen bonds between helical turns to stabilize each helix and thereby the dimer. Molecular dynamics simulations of an oligomer constructed with wall-forming core and transmembrane α9 dimers linked by flexible α6-α7-α8 bridges in a mitochondrial lipid bilayer generate an atomic resolution model for a stable Bax pore capable to release cytochrome C. Thus, we made an important step toward elucidating molecular mechanisms of apoptotic Bax perforation.

## Introduction

Bax is one of the two canonical pore-forming proteins in the Bcl-2 family that permeabilize the mitochondrial outer membrane (MOM) releasing mitochondrial proteins including cytochrome C to induce apoptotic death of over stressed or damaged cells; for reviews, see (Cosentino & Garcia-Saez, 2017; Kale *et al*, 2018b; Luna-Vargas & Chipuk, 2016; Moldoveanu & Czabotar, 2020; Spitz & Gavathiotis, 2022; Walensky, 2019). In healthy cells, Bax is mostly a soluble protein in the cytoplasm. The Bax protein structure determined by solution NMR is in a globular conformation with one core hydrophobic helix (α5) wrapped by seven amphipathic helices (Suzuki *et al*, 2000). In addition to the α5-α6 helical hairpin that is critical to the pore formation similar to its equivalents in the bacterial α-helical pore-forming proteins (Dal Peraro & van der Goot, 2016), the α9 is probably the most important helix that has the following demonstrated or proposed functions. In the soluble Bax conformation, it is an auto-inhibitory domain since it covers a binding groove for the BH3 proteins that activate Bax. This groove is canonical since it is shared among other Bcl-2 family proteins including the other pore-forming protein Bak, and the antiapoptotic proteins that bind to BH3 proteins to prevent them from activating Bax and Bak (Dlugosz *et al*, 2006). The antiapoptotic proteins also bind to the BH3 domain of active Bax or Bak such that these active molecules cannot use the exposed BH3 domain to activate other Bax and/or Bak (Cosentino *et al*, 2022; Iyer *et al*, 2020; Ruffolo & Shore, 2003; Tan *et al*, 2006). Moreover, the binding of antiapoptotic proteins to active Bax and Bak inhibits their homo-oligomerization that is required for their pore formation in the MOM (Ding *et al*, 2014; Ding *et al*, 2010). The mutually exclusive binding of the α9 helix of inactive Bax and of the α-helical BH3 domain of a BH3 protein or an active Bax to the same canonical groove provides a mechanism for the activation or autoactivation of Bax, respectively. Thus, the binding of the BH3 protein competitively displaces the α9 helix from the groove so it can function as a tail-anchor to target the Bax protein to the MOM whereas the empty groove in one active Bax can bind the exposed BH3 domain in other active Bax, and vice versa, forming a Bax dimer that is a building block for an oligomer (Dewson *et al*, 2012; Zhang *et al*, 2016; Zhang *et al*, 2010).

A second activation mechanism was proposed because some BH3 proteins can bind to a second site in the soluble Bax, which is located on the opposite side of the canonical groove (Dengler *et al*, 2019; Gavathiotis *et al*, 2008). Since this binding interaction triggers Bax conformational changes including releasing the α9 helix from the canonical groove, the binding site was termed the trigger pocket (Gavathiotis *et al*, 2010). Intriguingly, the trigger pocket of one Bax molecule is occluded by the α9 of the other Bax molecule as shown by the crystal structure of a soluble Bax dimer (Garner *et al*, 2016). Although the biological implication of this auto-inhibitory Bax dimer structure has been argued (Vogel *et al*, 2012), it nonetheless suggests another inhibitory function to the α9 helix, which instead of inhibiting the same Bax molecule *in cis* inhibits a different Bax molecule *in trans*.

Other than the auto-inhibitory and tail-anchor functions, the α9 helix has a third function based on the crosslinking data showing that it can interact with another α9 helix in another Bax that is also activated and targeted to the MOM (Zhang *et al*., 2016). Thus, the dimerization of the α9 helices mediates the higher order oligomerization of Bax molecules that have already dimerized via a reciprocal BH3-in-groove interaction, which was revealed by a crystal structure of a truncate Bax that only contain the core region (α2 to α5 helices) (Czabotar *et al*, 2013). The α9 dimerization was proposed to expand the Bax pore since larger Bax oligomers would form larger pores in membranes and a mutation that inhibited the α9 dimerization also inhibited the formation of the large but not the small pores (Zhang *et al*., 2016). However, a later study has challenged this proposal by showing that the same mutation shortens the residence time of Bax at the mitochondria in healthy cells and hence increases the threshold of apoptotic induction in stressed cells (Kuwana *et al*, 2020).

Even through many functions have been assigned to the α9 helix of Bax protein and some have been confirmed, there are many unknowns. The first unknown is the structure of the α9 dimer. Although several computational α9 dimer models of different configurations and stabilities in membranes have been constructed based on crosslinking data (Liao *et al*, 2016; Zhang *et al*., 2016), a high-resolution α9 dimer structure in any membrane-like environment has never been determined. Without the high-resolution structure, the function of the α9 dimer cannot be thoroughly tested by precise mutations that would disrupt the dimer structure or its interaction with the membrane.

The second unknown is the function of the α9 dimer in the oligomeric Bax pore assembly. Our recent paper (Lv *et al*, 2021) shows that an amphipathic α2-5 dimer forms part of the Bax pore wall that separates the hydrophobic lipid bilayer from the aqueous conduit. Since one α2-5 dimer is not large enough to form a complete wall that encircles a pore that can release the smallest apoptotic mitochondrial protein, cytochrome C, multiple α2-5 dimers are required to complete the pore wall. How multiple α2-5 dimers are linked to form the oligomeric pore is not known. The α9 is an obvious candidate for forming the interdimer linkage because it can dimerize with another α9.

The third unknown is the location of the α9 helix in the Bax pore. Both in plane and clamp models put it as a transmembrane helix deeply buried in the lipid bilayer (Bleicken *et al*, 2014; Westphal *et al*, 2014). However, the α9 helix is amphipathic so its hydrophobic side can cover the hydrophobic BH3 binding groove in the soluble Bax protein while leaving its hydrophilic side on the protein surface to engage the aqueous cytoplasmic environment. Even when it is in the membrane-bound Bax, it is not entirely buried in the membrane since some residues in the α9 helix are partially exposed to a water-soluble and charged labeling agent (Zhang *et al*., 2016). Quenching of fluorophores attached to residues in a9 by quenchers attached to different atoms in phospholipids and thus located at different depths in the bilayer suggests that a9 is either transmembrane or shallowly buried in the membrane (Bleicken *et al*, 2018; Kyrychenko *et al*, 2022). Release of fluorescent dyes from liposomes by α9 peptides suggests that these amphipathic peptides can form an aqueous conduit in the nonpolar lipid bilayer with some residues exposed to water (Garg *et al*, 2013).

To further determine the structure-function of Bax α9, in particular, when it is in a membrane, we first determined a high-resolution structure of the α9 in a lysolipid micelle using nuclear magnetic resonance (NMR) spectroscopy, which is a typical helical dimer structure glued by hydrophobic forces across the interface with the closely packed GxxxA motifs in the core. Interestingly, all the polar residues are exposed on the dimer surface. In addition to the mainchain-mainchain hydrogen bonds that are common to all α-helical structures, some of the polar residues also form rare sidechain-mainchain hydrogen bonds. These bifurcated hydrogen bonds are expected to further stabilize the helical structure of α9, which in turn may further stabilize the helical dimer. Mutations of both interfacial hydrophobic residues that disrupted the α9 dimer and the polar residues that eliminated the bifurcated hydrogen bonds reduced Bax pore formation. The defects caused by these α9 mutations in Bax pore formation were additive. Interestingly, these defects were also additive to the defect caused by a mutation in α4 that reduces the core dimer interaction with membranes (Lv *et al*., 2021). Therefore, stable dimerization of stable α9 helices in the mitochondrial membrane provides an important structural linkage between the pore-lining α2-5 dimers to facilitate the oligomeric Bax pore assembly. This conclusion was further supported by molecular dynamics (MD) simulations of a Bax oligomer that forms an aqueous pore in a model mitochondrial lipid bilayer, large enough to release cytochrome C.

## Results

### Structure determination of Bax α9 in micelles

To investigate the structure of Bax α9 in membrane mimic systems, we applied the NMR technology. We systematically tested different fusion tags and different detergent micelles and lipid bicelles for expression, purification and stability of the amphipathic and short α9 peptide (Bax residues Gly^166^ to Gly^192^). We were able to purify a Glu_3-_tagged α9 peptide and reconstitute it into dodecylphosphocholine (DPC) micelles to generate samples that were sufficiently soluble for structure determination (Fig. S1A). Chemical crosslinking experiments show that the peptides dimerize in the micelles (Fig. S1B).

A series of triple resonance experiments including HNCO, HNCACO, HNCA, HNCOCA, HNCACB and nnNOE were performed to assign the backbone amide resonances to the Glu_3_-α9 peptides in the micelles (Fig. 1A). The intramolecular nuclear Overhauser enhancement (NOEs) were collected from the isotopically labeled peptides in the micelle to assign the methyl group resonances (Fig. 1B). The intermolecular NOEs were measured between the [^15^N, ^2^H]-labeled and the [^13^C]-labeled peptides (Fig. 1C). Using 248 local and 48 long-range distance restraints derived from NOE measurements we determined the structures of an α9 ensemble in the micelles (Fig. 1D; Table 1).

**Figure 1.**
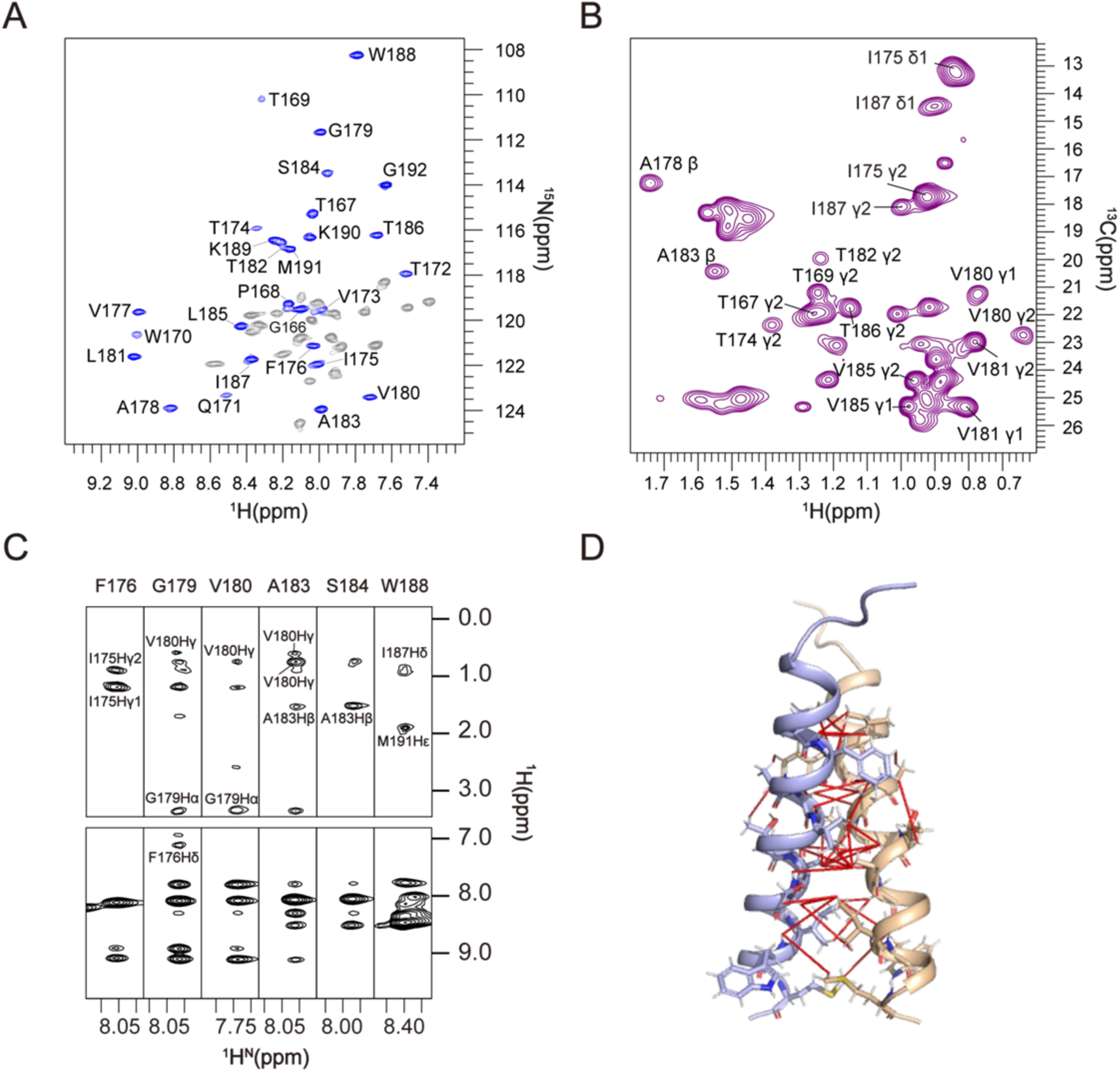
NMR spectra and intermonomer NOEs of Bax (α9) in micelles. **A**. Two-dimensional ^1^H-^15^N TROSY-HSQC spectrum of micelle-reconstituted Bax (α9) samples recorded at 600 MHz. The assigned backbone amide resonances of Bax (α9) (blue) and backbone amide resonances of E3 tag (grey) are shown. **B.** The 2D ^1^H-^13^C HSQC spectrum of Bax (α9) in deuterated DPC micelles recorded at 900 MHz with the assigned methyl group resonances. **C.** Intermonomer NOE strips from the mixed isotope labeled Bax (α9). The crosspeaks in the aliphatic regions are intermonomer NOEs between the backbone amide and the sidechain methyl protons. The spectrum was recorded with NOE mixing time of 200 ms at ^1^H frequency of 900 MHz. The sample used consists of 50% (^15^N, ^2^H)-labeled protein and 50% (^13^C)-labeled protein reconstituted in DPC micelles, in which the acyl chains of DPC were deuterated. **D.** Ribbon representation of the dimer structure with the NOE-derived intermonomer restraints (red lines).

**Table 1.** NMR and Refinement Statistics for Bax (α9) Structures.

|  |  |
| --- | --- |
| **NMR distance and dihedral constraints** |  |
| Distance constraints from NOE | 296 |
| Intra-chain NOEs | 248 |
| Inter-chain NOEs | 48 |
| Total dihedral angle restraints | 92 |
| $\Phi$ (TALOS) | 46 |
| $\Psi$(TALOS) | 46 |
| **Structure statistics<sup>a</sup>** |  |
| Violations (mean ± SD) | |
| Distance constraints ($\text{\AA}$) | 0.089±0.010 |
| Dihedral angle constraints ($^\circ$) | 0.555±0.025 |
| Deviations from idealized geometry |  |
| Bond lengths ($\text{\AA}$) | 0.006±0.000 |
| Bond angles ($^\circ$) | 0.191±0.032 |
| Impropers ($^\circ$) | 0.367±0.023 |
| Average pairwise RMSD ($\text{\AA}$) <sup>b</sup> | |
| Heavy atoms | 1.178 |
| Backbone | 0.672 |
| PDB ID | 6L95 |

<sup>a</sup> Statistics are calculated and averaged over an ensemble of the 15 lowest energy structures of the 200 calculated structures.
<sup>b</sup> The precision of the atomic coordinates is defined as the average RMSD between the 15 lowest energy structures and their mean coordinates.

The 15 lowest-energy α9 dimer structures (PDB ID: 6L95) out of 200 calculated structures converged to a root mean squared deviation (RMSD) of 0.672 Å and 1.178 Å for backbone and all heavy atoms, respectively (Fig. 2A; Table 1). In these high-resolution structures, the α9 helices form an intersected parallel dimer utilizing a classical transmembrane dimerization “Gly^179^-xxx-Ala^183^” motif (Fig. 2B). Four nonpolar residues, Phe^176^, Gly^179^, Val^180^ and Ala^183^, in one protomer interact with their counterparts in the other protomer forming the dimer core, which is extended by additional hydrophobic interactions on the N-terminal side between Ile^175^ and Phe^176^ in one protomer and Phe^176^’ and Ile^175^’ in the other protomer, respectively, and on the C-terminal side between Ile^187^ and Ile^187^’ (Fig. 2B). The resulting dimer interface extends along the entire length of the helix with a total contact surface area of 56.9 ± 4.7 Å^2^ as determined by PISA program. The dimer is approximately symmetrical and the C-termini of the interacting helices show a broader separation than the N-termini with an angle ∼27° ± 4° between the helical axes. A thio-specific nitroxide spin label, MTSL, was introduced into a few Glu_3_-α9 peptides, each with a residue replaced by a single cysteine, to collect paramagnetic resonance enhancement (PRE) data (Fig. S2A). The inter-monomer distance restraints derived from the PRE data are consistent with the structural model for the helical dimer (Fig. S2B).

**Figure 2.**
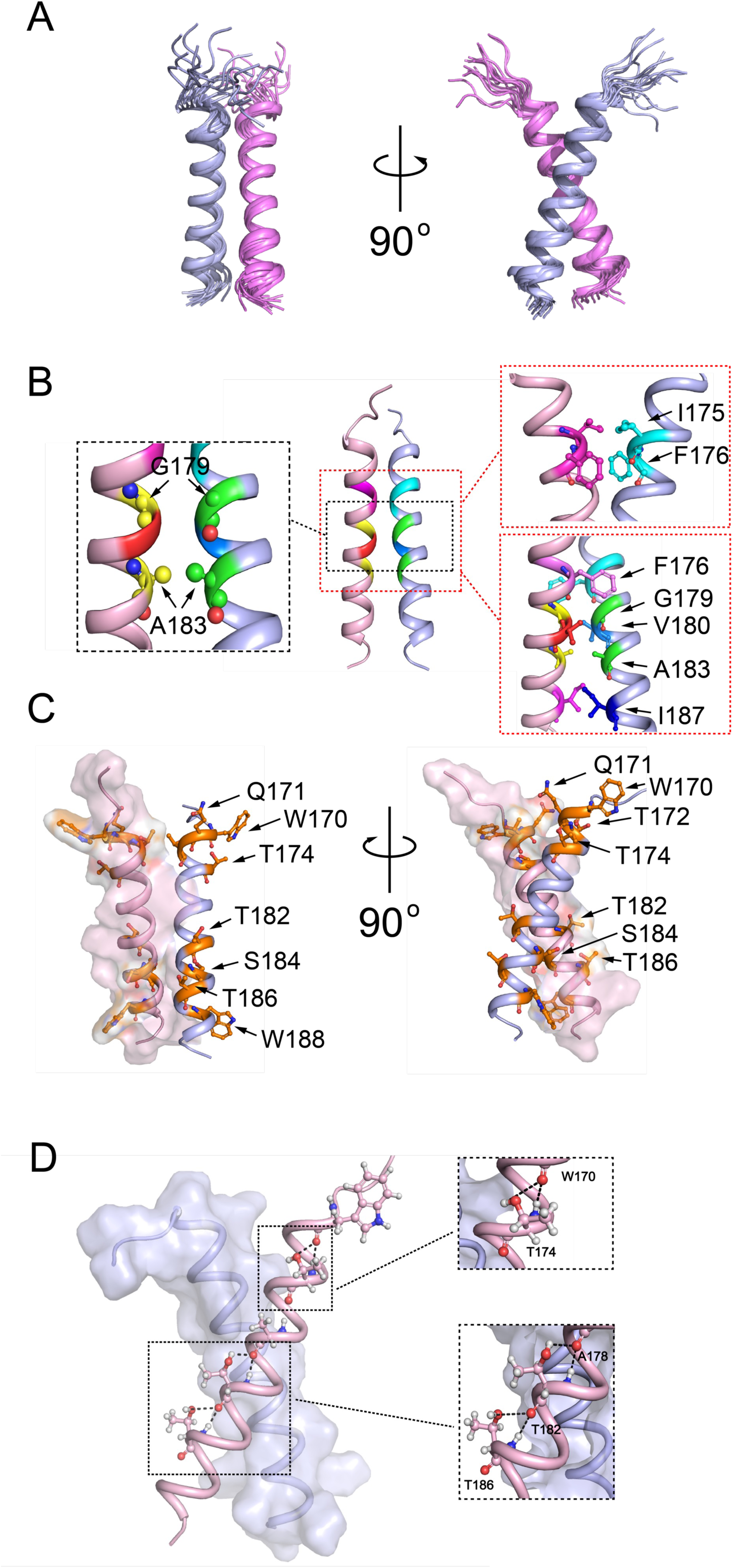
Structures of Bax α9 dimer in micelles. **A**. Ensemble of 15 low-energy structures calculated using NMR restraints summarized in Table 1. **B.** Ribbon representations of α9 dimer showing the interface with interacting GxxxA motifs and nonpolar residues enlarged in the left and right boxes, respectively. **C.** Ribbon and surface representations of a9 dimer showing polar residues on the surface between two nonpolar tryptophan residues. **D.** Ribbon and surface representations of a9 dimer showing potential bifurcated hydrogen bonds between T174, T182, T186 and T170, A178, T182, respectively.

In contrast to the α9 dimer interface that is formed by interacting nonpolar residues, the dimer surface contains many polar residues, including Gln^171^, Thr^172^, Thr^174^, Thr^182^, Ser^184^, and Thr^186^ (Fig. 2C). Some of these polar residues would be buried in the hydrophobic core of the lipid bilayer like their neighboring nonpolar residues such as Val^173^, Val^177^, Ala^178^, Leu^181^, and Leu^185^ if the α9 helices are transmembrane with Trp^170^ and Trp^188^ located in the polar lipid headgroup regions. According to an extensive survey of hydrogen bond geometries in different protein structures (references), the sidechain hydroxyls of Thr^174^, Thr^182^ and Thr^186^ have a high probability to form hydrogen bonds with the mainchain carbonyls of Trp^170^, Ala^178^ and Thr^182^, respectively, in the next helical turn (Fig. 2D; Fig. S3). These rare sidechain-mainchain hydrogen bonds together with the mainchain-mainchain hydrogen bonds between Trp^170^, Ala^178^, Thr^182^ and Trp^170^, Ala^178^, Thr^182^, respectively, common to all α-helical structures result in three bifurcated hydrogen bonds (Fig. 2D; Fig. S3) that would further stabilize the helical structure of α9 (Brielle & Arkin, 2020; Eswar & Ramakrishnan, 2000; Feldblum & Arkin, 2014), which in turn would further stabilize the helical dimer.

### The α9 dimerization plays an important role in the oligomeric Bax pore formation

Our previous crosslinking study has demonstrated that the α9 helices in active Bax proteins are in close proximity with each other such that the Bax proteins in the mitochondrial membrane can be crosslinked by a disulfide via the single cysteines placed along the helices (Zhang 2016 EMBO J). The atomic resolution structure of the α9 dimer we obtained here fits some of the published crosslinking data. In particular, in the α9 dimer structure, Ile^175^, Gly^179^, Ala^183^, and Ile^187^ in one protomer is in a disulfide-linkable distance with their counterparts in the other protomer (Fig. 3A). Indeed, when each of these residues was replaced by a cysteine, the resulting single-Cys Bax protein yielded a disulfide-linked dimer in the mitochondrial membrane after activation (Zhang 2016 EMBO J). In particular, the disulfide crosslinking of the Bax A183C mutant was the strongest, allowing us to monitor the α9 dimerization between active Bax proteins in the mitochondria robustly.

**Figure 3.**
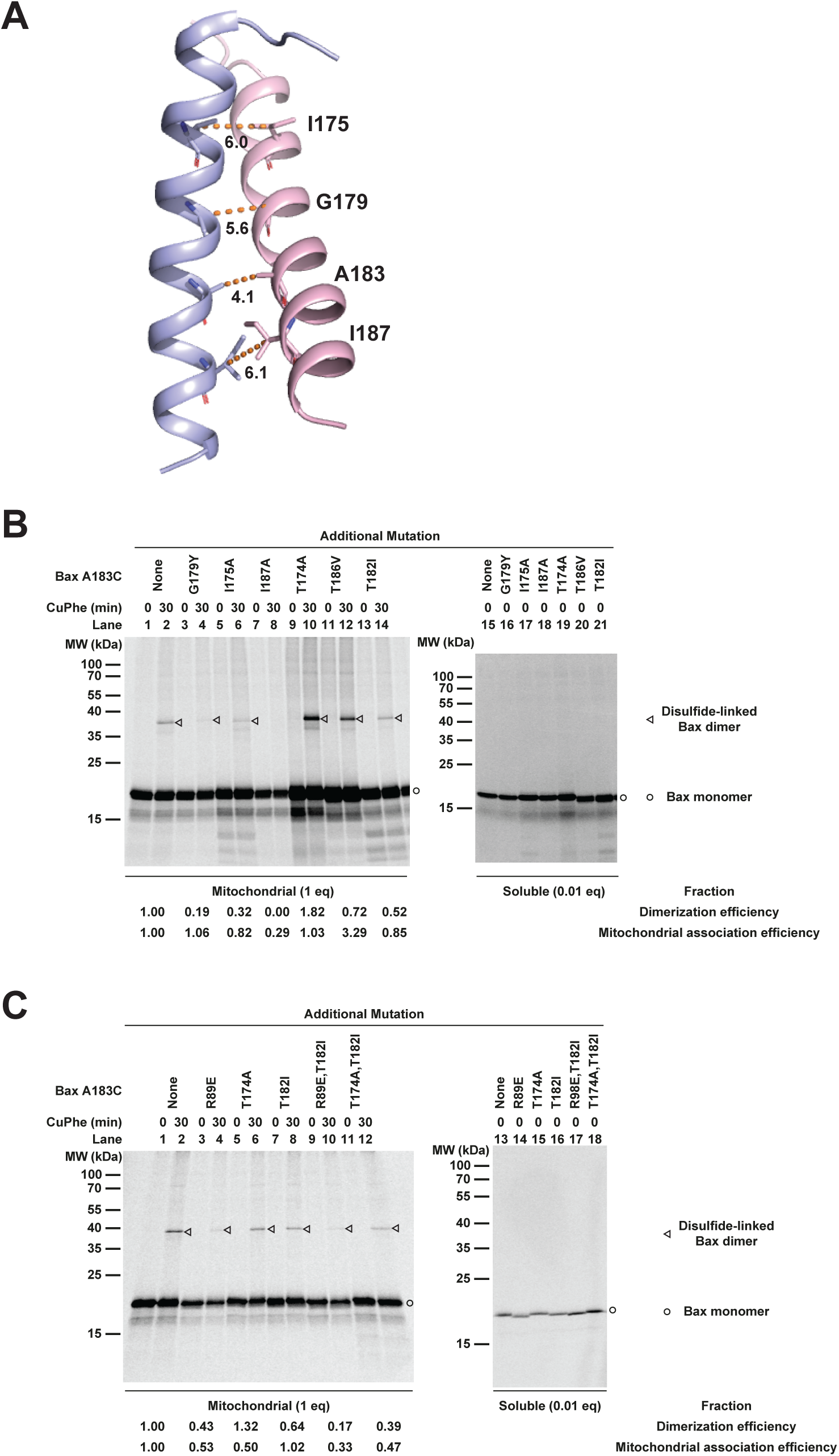
Structure-based mutations inhibit mitochondrial Bax dimerization via α9 helix. **A.** A ribbon diagram of the NMR structure of α9 dimer. The residue pairs that were replaced by Cys pairs for disulfide crosslinking described here and previously (Zhang *et al*., 2016) are shown as sticks with the Cβ carbons of the I175, A183 or I187 pair or the Cα carbons of the G179 pair linked by dashed lines. **B-C**. [^35^S]Met-labeled Bax proteins with a Cys replacing Ala^183^ (A183C) and additional mutations, if indicated, to disrupt the α9 or α2-α5 or both dimer interactions were synthesized, activated by Bax BH3 peptide, and targeted to the mitochondria lacking endogenous Bax and Bak proteins. The mitochondria-bound proteins were oxidized by CuPhe for 30 min to induce disulfide crosslinking of the two protomers via the A183C pair in the α9 dimer interface. The radioactive crosslinked Bax dimer was then separated from the monomer by non-reducing SDS-PAGE and detected by phosphor-imaging. The representative data from n = 3 or more independent experiments are shown. The relative dimer:monomer ratio shown at the bottom of the phosphor-images was determined by the intensities of the dimer and monomer bands of the A183C mutant with the additional mutation, normalized to that of the A183C mutant without the additional mutation. The mitochondrial association efficiency of each Bax mutant was determined by the summed intensity of the monomer and dimer bands in the mitochondrial fraction that was divided and treated with CuPhe for 0 or 30 min (I_M_), the intensity of the monomer band in the corresponding soluble fraction that was divided by 0.01 to adjust the inequivalent loading between the mitochondrial (1 equivalent (eq)) and soluble (0.01 eq) fractions (I_S_), and the following equation, E = I_M_/(I_M_ + I_S_). The relative mitochondrial association efficiency of each Bax A183C mutant with the additional mutation (E_R_) to that of the Bax A183C mutant without the additional mutation was obtained by dividing the E of the former with the E of the latter and shown at the bottom of the phosphor-images.

To determine the importance of the α9 dimerization in Bax pore formation, we first performed structure-guided mutagenesis to disrupt the dimer. Based on the structure, changing the small Gly^179^ to a bulky residue such as tyrosine would generate steric clashes in the dimer interface thereby disrupting the dimer. Indeed, we found that the G179Y mutation inhibited the α9 dimerization between active mitochondrial Bax proteins as detected by the A183C disulfide crosslinking (Fig. 3B, compare triangle-indicated bands in lanes 2 and 4). In contrast to the closely packed small glycine residues, the bulky residues such as Ile^175^ and Ile^187^ use their extended side chains to make favorable van der Waals contacts to further stabilize the α9 dimer. Mutating these bulky residues to smaller residues such as alanine would eliminate these favorable interactions thereby destabilizing the dimer. Accordingly, the I175A or I187A mutation reduced the A183C crosslinking and hence weakened the α9 dimerization (Fig. 3B, the triangle-indicated band in lane 6 is weaker than that in lane 2, and not detectable in lane 8).

The I175A or I187A mutation would decrease the hydrophobicity of α9 according to five hydrophobic scales (Kyte-Doolittle(Kyte & Doolittle, 1982), Wimley-White(Wimley & White, 1996), Hessa-Heijne(Hessa *et al*, 2005), Moon-Fleming(Moon & Fleming, 2011), and Zhao-London(Zhao & London, 2006)), whereas the G179Y mutation would increase the hydrophobicity according to four of the five scales (Wimley-White, Hessa-Heijne, Moon-Fleming, and Zhao-London) (Supplementary Table 1 and 2). Because α9 functions as a transmembrane domain that anchors the active Bax protein in the MOM, alteration of its hydrophobicity may alter Bax association with the mitochondria. Thus, we determined the mitochondrial association efficiency of the Bax A183C proteins without and with one of the dimer disruptive mutations. As expected, the I175A or I187A mutation decreased the mitochondrial association efficiency, whereas the G179Y mutation slightly increased the efficiency (Fig. 3B).

Next, we assessed the impact of these mutations on tBid-induced Bax pore formation in liposomal membranes that contain typical mitochondrial lipids by monitoring the release of fluorescent dye-labeled dextrans of 10 kDa whose hydrodynamic size is similar to that of cytochrome C. As shown in Figure 4A, the I175A mutation reduced the pore formation greatly such that the normalized fractions of fluorescent dextran release by the mutant were significantly lower than that by the wild-type from 7 minutes to the end of the 122-minute time course (from multiple unpaired t tests performed with the data sets of the wild-type (n = 4) and mutant (n = 3), P values ≤ 0.01 for differences of the means from 7 to 122 minutes). By the end of the time course ∼35% less dyes were released by the mutant than the wild-type. In comparison, the fractions of the dye release by the I187A mutant were not significantly different from that by the wild-type throughout the time course (P values ≥ 0.11). However, at a lower concentration, this mutant released significantly less dyes than the wild-type (Fig. S4A, P values ≤ 0.01 from 7 to 122 minutes). Surprisingly, the G179Y mutation significantly increased the fractions of dye release throughout the time course (P values ≤ 0.04), and by the end ∼37% more dyes were released by the mutant than the wild-type. The increase of the pore-forming activity by the G179Y mutation could be explained by the fact that this mutation auto-activated Bax in the absence of tBid resulting in ∼33% more dextran release than the wild-type by the end of the time course (Fig. S4B-D; P values ≤ 0.04 from 22 to 122 minutes).

**Figure 4.**
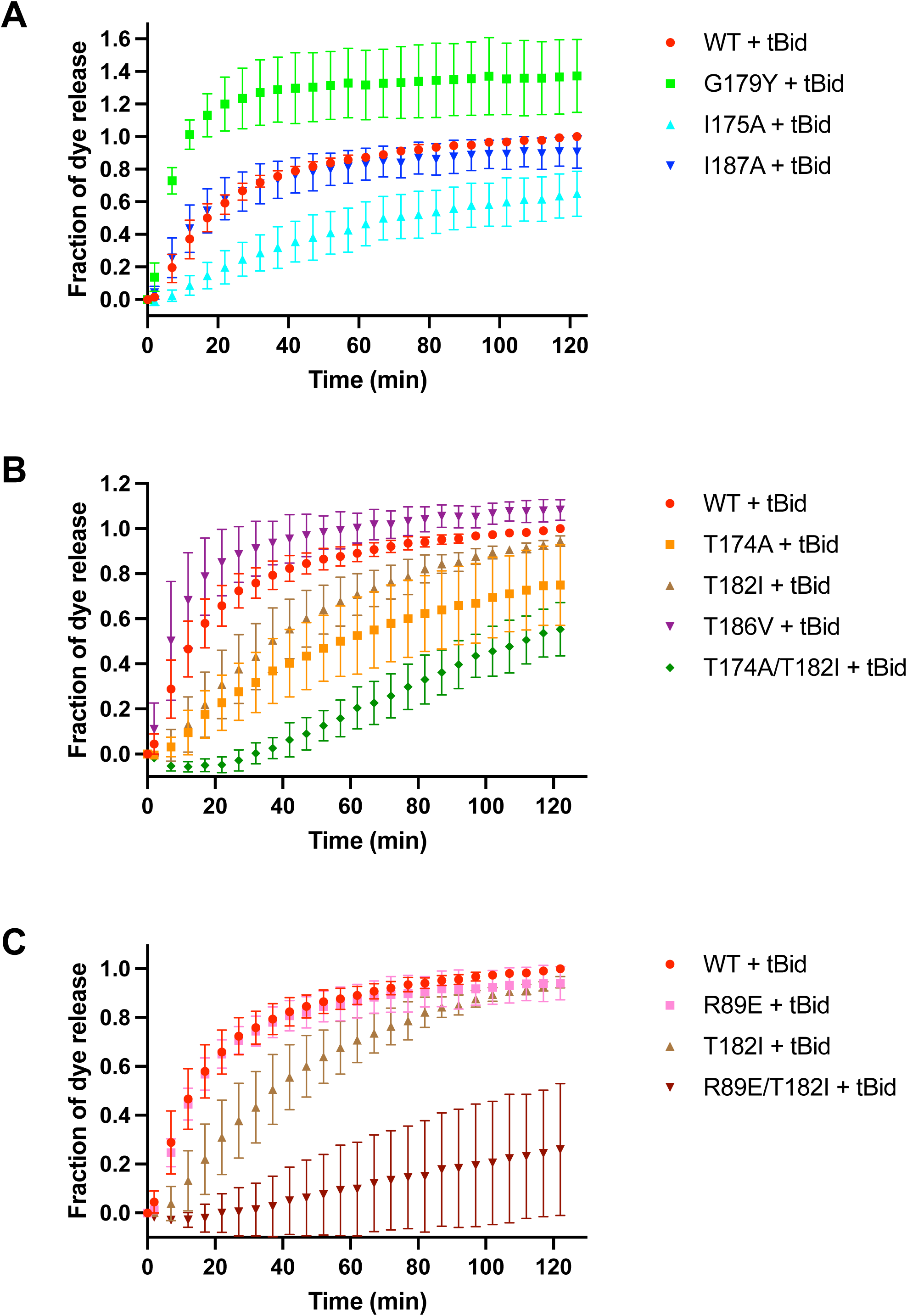
Structure-based mutations inhibit liposomal membrane permeabilization by Bax. Fluorescent dye release from 50 μM (total lipids) mitochondria-mimic liposomes by 100 nM wild-type (WT) or mutant Bax in the presence of 15 nM tBid was measured by quenching the fluorescence of the released dyes by the dye-specific antibodies outside the liposomes during a time course. The fraction of dye release was normalized to that by detergent, which was further normalized to the fraction of dye release by WT Bax and tBid at the end time point, The data shown were means (symbols) and standard deviations (error bars) obtained from n = 4 or 3 independent replicates for Bax WT or mutants, respectively.

To determine whether the mutations also alter the Bax pore formation in the mitochondrial membranes, we used the *bax* and *bak* double knockout baby mouse kidney (BMK) cells whose plasma membranes are permeabilized by a low concentration of digitonin so that purified Bax protein and its activator cBid can be delivered to the mitochondria inside the cells. Since the low concentration of digitonin barely permeabilizes the mitochondrial membranes, most of the mitochondria in the permeabilized cells have intact membranes and fluorescent mCherry protein expressed in the intermembrane space due to the mitochondrial targeting sequence from Smac protein fused to the mCherry protein. Thus, the Bax pore formation in the MOM can be monitored by measuring the release of the Smac-mCherry protein from the mitochondria. The fraction of Smac-mCherry release was measured before the addition of Bax and cBid and then at several time points after the addition, and normalized to that by the wild-type Bax and cBid at 60 min. As shown in Figure 5A, the fraction of baseline release measured before the Bax and cBid addition is ∼0.2, which did not increase after 60-min in the absence of Bax and cBid (Fig. S5), suggesting that most of the mitochondrial membranes in the permeable cells remain intact during the incubation as expected. Addition of Bax alone, either the wild-type or each of the three mutants, or cBid alone did not increased the release above the baseline (Fig. S5). In the presence of cBid, the wild-type Bax increased the fraction of release gradually, reaching 0.78 at 40 min, and plateaued at ∼1.00 after 60 min (Fig. 5A). In contrast, the I175A or I187A mutant did not increase the fraction of release significantly, only reaching 0.24 or 0.27, respectively, at 60 min. However, the G179Y mutant increased the fraction of release more than the wild-type did, reaching 1.00 at 40 min, and 1.11 at 60 min. Thus, the I175A and I187A mutations significantly reduced the Bax pore-forming activity in the MOM (from multiple unpaired t tests performed with the data sets of the wild-type and mutants at 40 and 60 min, P values ≤ 0.00005), whereas the G179Y mutation significantly increased the activity (with data sets at 20, 40 and 60 min, P values ≤ 0.03).

**Figure 5.**
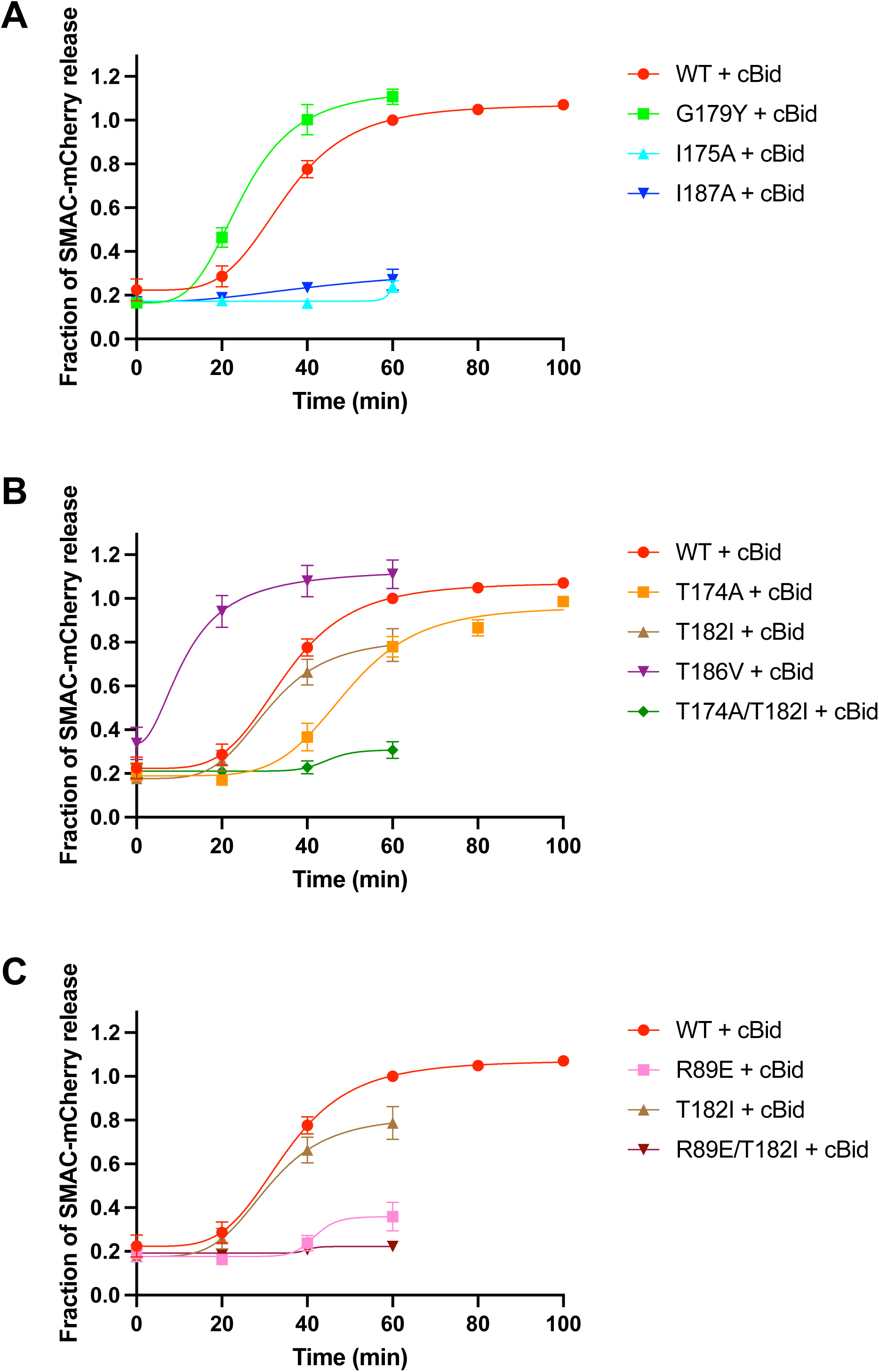
Structure-based mutations inhibit mitochondrial membrane permeabilization by Bax. Digitonin permeabilized BMK Bax^-/-^/Bak^-/-^ cells expressing SMAC-mCherry in the mitochondria intermembrane space (1,300,000 total cells) were incubated with 50 nM of WT or mutant Bax and 5 nM cBid for the indicated time at 37°C. The samples were separated into supernatant and pellet fractions by na centrifugation. Both fractions were then further solubilized by Triton X-100. The soluble materials from both fractions were collected after a centrifugation and their mCherry fluorescence spectra were measured. SMAC-mCherry release was calculated as the fraction of total (supernatant + pellet) mCherry fluorescence coming from the supernatant fraction. The data was normalized to the fraction of SMAC-mCherry release by WT Bax and cBid at 60 min. Each symbol represents the mean SMAC-mCherry release from n = 3 or more independent replicates with the error bar representing standard error of the mean. Each line represents a nonlinear square fit of the data with the [Agonist] vs. response - Variable slope (four parameters) equation in GraphPad Prism 10.5.0.

The effects of the mutations on Bax pore formation in the liposomal and mitochondrial membranes are qualitatively consistent (Fig. 4A vs. Fig. 5A), except that the G179Y mutant is autoactive in the liposomal but not mitochondrial membrane (Fig. S4A vs. Fig. S5). Overall, the functional data from the structure-guided mutagenesis experiments demonstrate that the α9 dimerization is important to the Bax pore formation in the mitochondrial membrane, particularly when less Bax proteins are associated with the mitochondria. However, gain of mitochondrial association and autoactivation of Bax can compensate the loss of α9 dimerization resulting in faster and more Bax pore formation.

### The bifurcated hydrogen bonds that are expected to stabilize the α9 helix contribute to oligomeric Bax pore formation

A novel feature revealed by the atomic resolution structure of the α9 dimer is the potential hydrogen bonds between the sidechain hydroxyls of Thr^174^, Thr^182^ and Thr^186^ and the mainchain carbonyls of Trp^170^, Ala^178^ and Thr^182^, respectively (Fig. 2D and S3). Since these hydrogen bonds as part of the bifurcated hydrogen bonds between adjacent helical turns may further stabilize the α9 helix in membranes, which in turn may further stabilize the α9 dimerization and Bax pore, we mutated each of the threonine residue to a nonpolar residue to eliminate one of the three sidechain-mainchain hydrogen bonds and measured the effect on the dimerization and pore formation. As shown in Figure 3B, the T182I or T186V mutation reduced, whereas the T174A mutation elevated, the Bax dimerization via α9. Although all the mutations are expected to increase the hydrophobicity of α9 (Supplementary Table 1 and 2), only the T186V increased the mitochondrial association (Fig. 3B). The T174A or T182I either did not change or rather decreased the association, respectively.

In the liposomal system, the T174A mutation decreased the Bax perforating activity more than the T182I, both significantly throughout the time course (P values ≤ 0.03 from 7 to 122 min), whereas the T186V significantly increased the activity after 62 min (P values ≤ 0.03) (Fig. 4B). The effects of these mutations in the mitochondrial system largely mirrored that in the liposomal system (Fig. 5B, T174A vs. wild-type, P values ≤ 0.05 at 20, 40, 60 and 80 min; T182I vs. wild-type, P value = 0.007 at 60 min), except that the T186V mutant was autoactive in the mitochondria (Fig S5, P value < 0.0001 at 60 min) but not in the liposomes (Fig. S4B, P values ≥ 0.19 throughout the time course). Notably, combination of the T174A and T182I exacerbated the negative effect of the individual mutations, further decreasing Bax dimerization via α9 in mitochondria (Fig. 3C, compare triangle-indicated bands in lanes 6, 8 and 12 and the derived dimerization efficiencies) while not altering Bax association with mitochondria. Consistently, the double mutant further decreased the perforating activity in both liposomes (Fig. 4B, P values ≤ 0.007 from 7 to 122 min) and mitochondria (Fig. 5B, P values ≤ 0.00005 at 40 and 60 min). Thus, the potential intrahelical sidechain to mainchain hydrogen bonds in α9 that most likely stabilize the helix contribute to the oligomeric Bax pore formation, in particular when the mitochondrial association of Bax is not increased and Bax is not autoactive.

### The antiparallel α2-5 dimer and the parallel α9 dimer are two critical parts for an oligomeric Bax pore assembly

Previously we have determined the NMR structure of an antiparallel dimer formed by Bax α2 to α5 helices in bicelles, and demonstrated this amphipathic dimer functions as part of the apoptotic pore wall in the mitochondrial membrane (Lv et al EMBOJ 2021). However, it was not known how multiple α2-5 dimers are linked together to encircle the aqueous pore. The α9 dimer structure we determined here raised a possibility for a parallel α9 dimer to link two antiparallel α2-5 dimers. Moreover, the α9 dimer fully embedded in a lipid bilayer could anchor the amphipathic α2-5 dimers in the wall that separates the nonpolar lipid bilayer from the aqueous pore. To determine whether the two dimers work together to form pores in membranes, we chose the R89E mutation in α4 that disrupts the polar interaction of α2-5 dimer with the negatively charged lipid headgroups in the bilayer, and partially reduces the Bax perforation (Lv et al EMBOJ 2021). We combined this mutation with the T182I mutation in α9 that also partially reduces the Bax perforation to see if their negative effects on the perforation are additive. As shown in Figure 4C, the T182I alone significantly reduced the Bax perforation in the liposomal membrane (P values ≤ 0.03 from 7 to 122 min), whereas the R89E alone did not (P values ≥ 0.13 throughout the time course). However, at a lower concentration, the R89E released significantly less dyes than the wild-type (Fig. S4A, P values ≤ 0.05 from 2 to 122 minutes). When combined with T182I, however, the R89E mutation further reduced the Bax perforation (P values ≤ 0.009 from 7 to 122 min). In the mitochondrial membrane, the reduction of Bax perforation by the R89E (P values ≤ 0.00001 at 40 and 60 min) was more than that by the T182I, and their combination further reduced the perforation (Fig. 5C, P values ≤ 0.00003 at 40 and 60 min). Consistently, the negative effects of the two mutations on Bax dimerization via α9 were also additive (Fig. 3C, compare triangle-indicated bands in lanes 4, 8 and 12 and the derived dimerization efficiencies). Therefore, by simultaneously disrupting the functions of two critical parts, the impact of the two mutations together on the overall Bax pore formation is more profound than that of each mutation alone.

### A pore model generated from molecular dynamics simulations of a Bax oligomer in a mitochondrial lipid bilayer

To generate a model for a Bax pore that can release cytochrome C, we first constructed a Bax oligomer with 16 α2 to α9 regions in a lipid bilayer comprising mitochondrial characteristic lipids. The adjacent α2-5 regions of the oligomer dimerize according to the previous NMR structure (PDB ID: 6L8V) and form a circling wall that separates the aqueous conduit from the lipid bilayer as we previously demonstrated (Lv *et al*., 2021). The adjacent α9 regions dimerize according to the current NMR structure (PDB ID: 6L95) and embeds in the lipid bilayer as we previously determined (Zhang *et al*., 2016). Each α2-5 region is linked to the downstream α9 region by the α6-8 region in an elongated conformation with each helix modeled after the corresponding helix in a soluble Bax structure (PDB ID: 1F16) previously determined by NMR (Suzuki *et al*., 2000). From this initial pore configuration surrounded by water molecules and ions (Fig. S6A), we performed all atom molecular dynamics (MD) simulations.

By the end of a 700-ns simulation and ten independent replicates, stable pore structures evolved in the lipid bilayer (Fig. 6A, and S6B). The eight amphipathic α2-5 dimer structures remain stable (RMSD < 2.0 Å compared with the initial structure for residues 48-124) with the polar α2-3 surface lining the aqueous conduit and the nonpolar α4-5 surface engaging the lipid bilayer. Some of the pores display a circular shape with a diameter of ∼60 or 70 Å (Fig. S6B, top view, middle or right diagram, respectively), while other an elliptical shape with a major axis of ∼100 Å and a minor axis of ∼40 Å (Fig. S6B, top view, left diagram). The relative orientations between adjacent α2-5 dimers are similar in the circular pores (a representative one is boxed in Fig. S7A, left diagram), whereas they are different in the elliptical pore (two representative ones are boxed in Fig. S7A, right diagram), suggesting that different modes of interactions may exist between some dimers.

**Figure 6.**
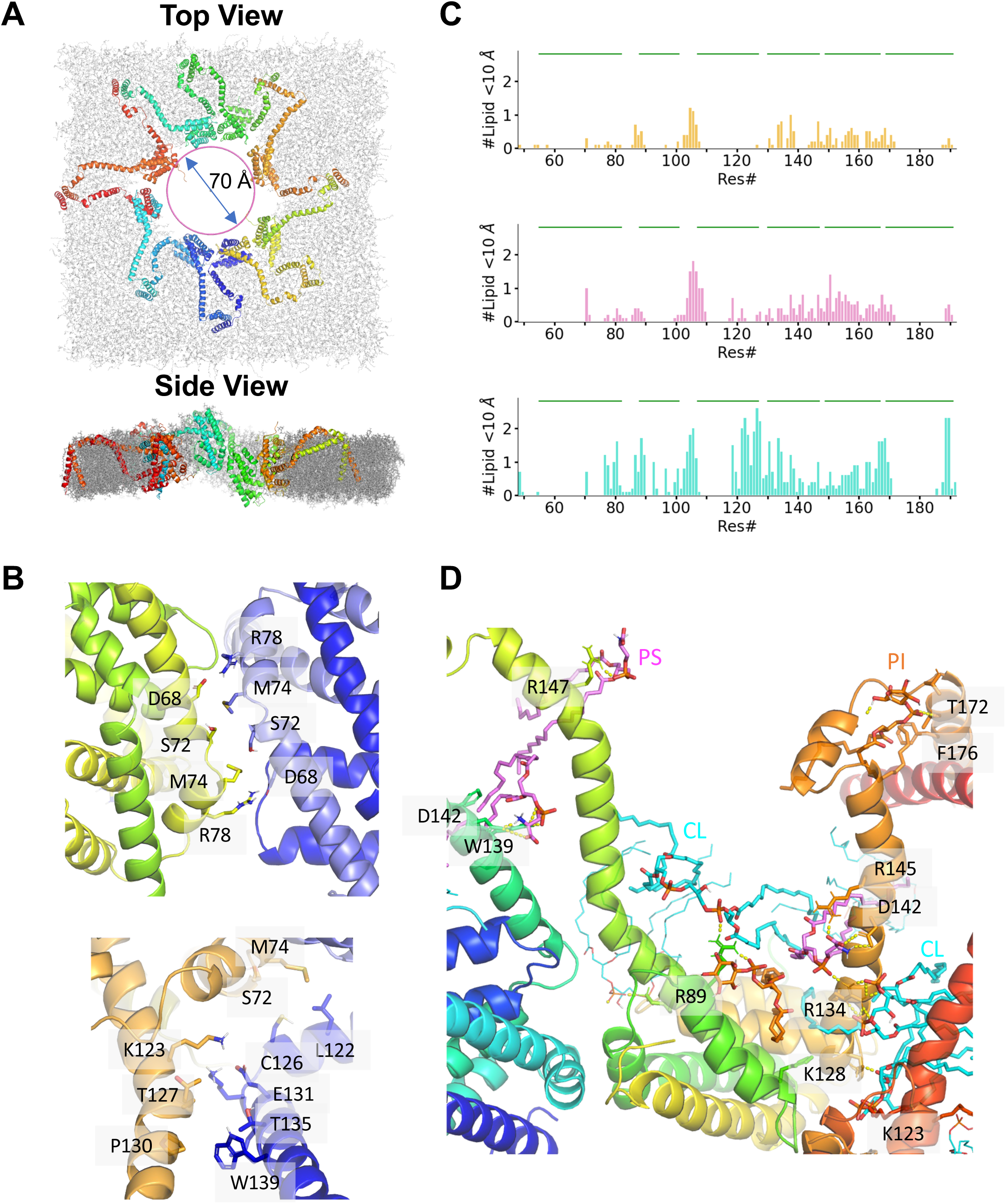
Oligomeric Bax pores in a model mitochondrial lipid bilayer revealed by molecular dynamics (MD) simulations. All-atom MD simulations of a pore assembled from 16 interacting Bax molecules each containing the α2 to α9 region in a model mitochondrial lipid bilayer were carried out with CHARMM36m force field for 700 ns. The resulting pores have the following features. **A.** Each Bax molecule interacts with the neighboring Bax molecules via a α2-5 dimerization and a α9 dimerization. Pores of ∼40-70 Å wide were formed at the end of ten independent simulations. **B.** Potential contacts between neighboring α2-5 dimers are illustrated by the indicated residues whose Cý atoms are within 6 Å. **C.** Potential protein-lipid contacts are indicated in the top, middle and bottom graph, respectively, by the number of PI, PS, and CL that are within 10 Å of Cα of Bax residues. **D.** Potential hydrogen bonds between the indicated Bax residues and PI, PS, and CL are indicated by yellow dashed lines.

Majority of the eight α9 dimers remain in the intersected conformation with the GxxxA motifs at a close distance (Fig. S7B, left diagram). These amphipathic α9 dimer structures remain stable (RMSD < 2.0 Å compared with the initial structure for residues 170-192) and fully embedded in the lipid bilayer. However, few α9 dimers change the conformation with less interactions between the GxxxA motifs, such as those in the elliptical pore (Fig. S7B, right diagram). In both α9 dimer conformations, the bifurcated H-bonds between W170 and T174, and between A178, T182 and T186 are persistent during the 700-ns simulations (Fig. S7B), most likely because they further stabilize the amphipathic helical dimers in the mitochondria-mimic lipid bilayer.

In contrast, some of the sixteen α6-8 regions are less stable resulting in heterogeneous structures in heterogeneous environments (RMSD ∼ 2.3 - 5.1 Å k compared with the initial structure, and between the final structures for residues 128-165). In particular, one α6 was merged with the upstream α5 and the downstream α7 forming an extended helix with kinks near the α5/α6 and α6/α7 boundaries (Fig. S8A, Chain B), while the other α6 was still connected to the α5 via a loop but merged with the downstream α7 with a kink in the boundary (Fig. S8A, Bax Chain C). Some of the α7 and α8 helices were unfolded partially while others were not (Fig. S8B, α7 in Chain J is shorter than α7 in Chain K, whereas α8 in Chain K is shorter than α8 in Chain J). Half of the α6 regions are partially embedded in the upper leaflet of the lipid bilayer presumably facing the cytosol (e.g., Fig. S8C left, α6 in Chain K), whereas the other half cross the lipid bilayer from the lower leaflet (presumably facing the mitochondrial intermembrane space) to the upper leaflet (e.g., Fig. S8C right, α6 in Chain A).

Thus, these oligomeric pores are formed by eight rigid wall blocks (α2-5 dimers) and eight rigid transmembrane anchors (α9 dimers) that are connected by sixteen flexible bridges (α6-8 regions) through different parts of the lipid bilayer. The sizes of the pores by the end of different simulations were varied ranging from 40 to 70 Å that are comparable to the diameter of a hydrated cytochrome C ∼60 Å (Fig. S6B), likely capable to release this smallest apoptogenic mitochondrial protein and close to what we estimated before (Lv *et al*., 2021).

In addition to the dimeric α2-5 and α9 regions, close contacts between other regions of neighboring Bax molecules around the pore were revealed by the computational pore models. For example, the α2 and α3 in a α2-5 dimer are close and antiparallel to the α2 and α3 in another α2-5 dimer in part of the pore wall (Fig. 6B top, D68, S72, M74 and R78 in one dimer are within 6 Å of R78, M74, S72, and D68 in the other dimer, respectively; Supplementary Table S3). In another part of the pore wall and bridge, the α5 and α6 in a Bax molecule are proximal to the α2, α5 and α6 in another Bax molecule (Fig. 6B bottom, L122, C126, E131, T135 and W139 in one molecule are within 6 Å of M74, S72, K123, T127 and P130 in the other molecule, respectively; Supplementary Table S3).

On the other hand, gaps between some of the protein wall blocks could also be seen in the computational models, and they were filled by lipids including phosphatidylcholine, phosphatidylethanolamine and cardiolipin (Fig. S9). Some of the lipids in the gaps are not in the typical bilayer configuration; instead, they rearranged toward micelle configuration with the polar heads pointed to the aqueous pore lumen and the nonpolar tails point to the nonpolar core of the bilayer (Fig. S9).

Specific lipids preferentially associated with specific Bax regions in the perimeter of the pore according to the computational models. Negatively charged phosphatidylinositol, phosphatidylserine and cardiolipin were found within 6 Å of the loop between α4 and α5 helices (Fig. 6C, top, middle and bottom graphs, respectively). They were also close to α6, α7 and α8 helices and the connecting loops. Moreover, cardiolipin, but not the other two phospholipids, was close to the C-terminal halves of α5 and α9. Overall, more cardiolipins were close to the pore-forming proteins than the other two phospholipids. Some of these negatively charged phospholipids formed hydrogen bonds with protein residues, especially the charged residues D71, R89, K119, K123, K129, R134, R145 and R147, the uncharged polar residues T127 and T169, and the nonpolar residue W107 (Fig. 6D; Fig. S10 and S11; Supplementary Table S4).

The lipid bilayer seemed to curve around the edge of Bax pores as observed by cryo-electron microscopy (Kuwana *et al*, 2016). Interestingly, curvature was seen in the lipid bilayer around the computational pores (Fig. 6A side view, S6B side view, and S6C). The curvature could be generated by uneven distribution of protein regions in the two leaflets of the lipid bilayer. As noted above (Fig. S8), some α6-8 regions cross the lipid bilayer from one leaflet to the other and along the way they increase the volume of one and then the other leaflet. In addition, the curvature could be caused by uneven distribution of protein wall blocks relative to the plane of the lipid bilayer (Fig. S6D). Thus, some α2-5 dimers are higher than the plane whereas others lower, and they may drag the associated lipids with them to create curvature.

In essence, our computational pore models not only suggest a minimal number of Bax proteins that can form an oligomeric pore large enough to release cytochrome C but additional contacts between these pore-forming proteins. More impressively the models also suggest contacts between phospholipids and Bax proteins that could be important for forming the protein-lipid pore as proposed before (Bleicken *et al*., 2014).

## Discussion

We made the following major discoveries in this structure and function study of the α9 dimerization between the mitochondrial membrane embedded perforating Bax proteins, whose impacts to the apoptosis and membrane protein fields were discussed below considering other important studies.

*(i) A high-resolution structure of a complex formed by two α9 peptides in lysolipid micelles determined by multifaceted solution NMR spectroscopy*. The interface of the α9 dimer is formed by intercalating nonpolar residues centered around the closely packed GxxxA motifs, also found in many other transmembrane helical dimers; for reviews, see (Curran & Engelman, 2003; Fink et al, 2012; Teese & Langosch, 2015). In contrast, the surface of the α9 dimer is amphipathic with nonpolar residues that are expected to interact with the nonpolar core of a lipid bilayer, and polar residues that are present in many other transmembrane helices. Because some of these polar residues would be surrounded by the nonpolar tails of lipids, the hydrogen bonding potential of their sidechain hydroxyls could not be fulfilled by water molecules or ions that are scarce in the nonpolar core of a lipid bilayer. An alternative solution would be for these polar residues to form bifurcated hydrogen bonds. Indeed, the high-resolution α9 dimer structure reveals potential hydrogen bonds between the sidechain hydroxyls of Thr^174^, Thr^182^ and Thr^186^ and the backbone carbonyls of Trp^170^, Ala^178^ and Thr^182^, respectively. In addition, the backbone carbonyls of Trp^170^, Ala^178^ and Thr^182^ still retain the potential to form hydrogen bonds with the backbone amides of Thr^174^, Thr^182^ and Thr^186^, respectively, thereby resulting in three bifurcated hydrogen bonds that would further stabilize the α9 dimer structure in the mitochondrial membrane.

In general, protein structures contain proper arrangements of H bonds. Many of H bonds involve a single donor and acceptor pair such as the regular H-bonding pattern between carbonyl oxygens and amide hydrogens four residues apart in α-helices. However, the hydrogen bonding potential of some acceptors leads to over coordination between two donors and one acceptor, resulting a bifurcated H bond. The strength of such bifurcated H bonds in a transmembrane α-helix was determined by using isotope-edited Fourier transform infrared (FTIR) spectral measurements and density functional theory (DFT) calculations (Feldblum & Arkin, 2014). The strength of the hydroxylic bifurcated H bond is -3.4 kcal/mol. In comparison the canonical H bond found in α-helices is -5,7 kcal/mol. The bifurcated H bond does not necessarily weaken the canonical H bond due to that it does not compete with it. The strength of all H bonds depends greatly on their surroundings, stronger in nonpolar and weaker in polar environments. A thermodynamic study of α helix-rich protein shows canonical H bonds can be stronger by up to 1.2 kcal/mol when they are sequestered in hydrophobic surroundings than when they are solvent exposed (Gao *et al*, 2009). The formation of bifurcated H bonds would also be favor in hydrophobic milieus, e.g., solvent inaccessible cores of water-soluble proteins and transmembrane domains of integral membrane proteins. The strength of the intrahelical bifurcated H bond is consistent with its prevalence in these hydrophobic environments. A survey conducted by Brielle and Arkin reveals 92% of all transmembrane helices having at least one non-canonical H bond form by a serine or threonine residue at position *i* whose hydroxyl sidechain H-bonds to an over-coordinated carbonyl oxygen at position *i-4*, *i-3* or *i* in the sequence (Brielle & Arkin, 2020). Isotope-edited FTIR spectroscopy and DFT calculations allowed them to determine the bond enthalpies, resulting in values that are up to 127% higher than that of a single canonical H bond. These strong H-bonds are expected to stabilize these polar residues in hydrophobic milieus while at same time providing them flexibility to adapt different configurations necessary for function.

In another survey, 19,835 polar residues from 250 non-homologous protein crystal structures were used to identify sidechain-mainchain (SC-MC) H bonds (Eswar & Ramakrishnan, 2000). The ratio of the number of SC-MC H bonds to the total number of polar residues is about 1 to 2. Close to 56% of these H bonds are formed by a SC acceptor/donor at *i* position and a MC donor/acceptor within a window of *i-5* to *i+5* position. These short-range H bonds for well-defined conformational motifs characterized by specific combinations of backbone and side chain torsion angles. The serine and threonine residues show the greatest preference in forming intrahelical H bonds between the atoms O^ψ^*_i_* and O*_i-4_*, and more than half such H bonds are found at the middle of α-helices rather than at their ends.

Many transmembrane helices contain serine and/or threonine residues whose sidechains for intrahelical H bonds with upstream carbonyl oxygens. The impact of threonine sidechain-mainchain H-bonding on the backbone dynamics of the amyloid precursor protein transmembrane helix was studied (Scharnagl *et al*, 2014). Circular dichroism spectroscopy and amide exchange experiments of transmembrane peptides reveal that mutating threonine enhances the flexibility of this helix. Molecular dynamics simulations show that the mutations reduce intrahelical amide H-bonding and H-bond lifetime. The removal of sidechain-mainchain H-bonding distorts the helix, which alters bending and rotation at a hinge connecting the dimerization region and the cleavage region that is processed by γ-secretase. The backbone dynamics of the substrate may profoundly affect how it is presented to the catalytic site of the enzyme. Changing this conformational flexibility may change the pattern of proteolytic processing, thereby explaining how the threonine mutations lead to early onset of Alzheimer’s disease.

The bifurcated H bonds can also be formed by other residues. The conformation of a polyglutamine tract of the androgen receptor associated with spinobulbar muscular atrophy depends on its length. This sequence folds into an α helical structure stabilized by non-canonical H bonds between glutamine sidechains and mainchain carbonyl groups and its helicity directly correlates with the tract length (Escobedo *et al*, 2019). These unusual H bonds are bifurcated with the canonical H bonds further stabilizing the α-helices, establishing the mechanistic basis of the disease.

Ser^184^ in the α9 of Bax is a target of a pro-survival kinase that regulates intracellular localization and apoptotic activity of Bax. Phosphorylation of Ser^184^ by Akt coverts the proapoptotic Bax to an antiapoptotic protein because the phosphorylation enables Bax binding to proapoptotic BH3-only proteins in the cytosol and prevents Bax insertion into mitochondria (Kale *et al*, 2018a). Akt is frequently activated in cancers resulting in increased phosphorylated antiapoptotic Bax that promotes resistance of cancer cells to inherent and drug-induced apoptosis. In all of 20 soluble Bax structures determined by NMR Ser^184^ is buried in the BH3-binding groove. For the phosphorylation to occur the soluble Bax must change conformation to expose Ser^184^ to Akt. As discussed later a fraction of Bax proteins in solution and cytosol could assume a different conformation that is different from the NMR structures, with the α9 swinging out of the groove and hence making Ser^184^ accessible to Akt. On one hand, the phosphorylation adds a negative charge to the Ser^184^, which could keep the α9 out of the hydrophobic groove, enabling the binding of BH3-only proteins to the groove. On the other hand, the addition of a negative charge to the α9 could keep it out of the mitochondrial membrane, disabling the targeting of Bax to mitochondria. Interestingly, in about a half of the 20 NMR structures of soluble Bax a bifurcated hydrogen bond could potentially be formed between Ser184 and Val180 since the sidechain hydroxyl and the backbone amide of Ser^184^ are proximal to the backbone carbonyl of Val^180^ (Suzuki *et al*., 2000). In contrast, only two NMR structures show a proximity of the sidechain hydroxyl of Ser^184^ to the sidechain carboxyl of Asp^98^. Thus, formation of the sidechain-mainchain hydrogen bond between Ser^184^ and Val^180^ in the α9 might compete with formation of the sidechain-sidechain hydrogen bond between Ser^184^ in the α9 and Asp^98^ in the α4. Switch of an interhelical to an intrahelical hydrogen bond might reduce the affinity of α9 to the groove thereby facilitating the exposure of both α9 and the groove to the cytosol for interaction with other proteins or membranes.

*(ii) A systematic assessment of the contribution of the α9 dimer structure to the oligomeric Bax pore assembly in the mitochondrial membrane*. We determined the effect of a series of structure-guided mutations on Bax oligomerization and perforation here and previously (Zhang et al., 2016). As expected, most of the mutations that reduce the nonpolar interactions in the α9 dimer interface reduced Bax dimerization via the α9 helices and Bax perforation in the model and native mitochondrial membranes. Most of the mutations that eliminate the bifurcated hydrogen bonds also reduced the Bax interaction and perforation, likely due to a reduction of the stability of a9 in the mitochondrial membrane as discussed above.

(*iii*) *A model for oligomeric Bax pore consists of α2-a5 dimers and α9 dimers.* Previous studies from us and others have demonstrated that the core regions of Bax containing α2 to α5 helices form an antiparallel dimer (Czabotar *et al*., 2013; Dewson *et al*., 2012; Lv *et al*., 2021; Zhang *et al*., 2016). The dimer surface formed by α4 and α5 helices can bind to a lipid bilayer through both nonpolar interactions with the lipid acyl chains and polar interactions with the lipid headgroups. The dimer surface formed by α2 and α3 helices is mostly polar and exposed to an aqueous milieu. Therefore, an amphipathic core dimer can form part of the Bax pore wall in the mitochondrial membrane. To complete the pore wall, multiple core dimers are required. Because biologically meaningful interactions between core dimers have not been detected, the multiple core dimers are presumably linked by other regions of Bax. The α9 dimer that we have structurally and functionally characterized can serve as a linker between the pore-lining core dimers. A combination of the R89E mutation that reduces the interaction of core dimer with the lipid bilayer and the T182I mutation that reduces the dimerization of α9 helices within the lipid bilayer inhibited the Bax perforation in the mitochondrial membrane more than each mutation alone could do (Fig. 4 and 5). This result has testified the important contributions of these two dimer structures to the oligomeric Bax pore assembly in the mitochondrial membrane, as illustrated in the computational model generated by MD simulations (Fig. 6).

(iv) *NMR structures of α2-5 dimer and α9 dimer and Bax pore model from MD simulations versus cryoEM structures of Bax pore.* While this manuscript is in preparation, the Shi and Huang laboratories published structures of Bax pores determined by cryoEM (Zhang *et al*, 2025). This milestone discovery in apoptosis field revealed overall structures of Bax pore formed by repeating units. Each repeating unit contains four Bax protomers, organized as a dimer of asymmetric dimers (Fig. S12A). The two protomers in an asymmetric dimer show different conformations. In type I protomer (e.g., the green chain) α6 and α7 merge into a single helix that is linked to α8 via a short loop, whereas in type II protomer (e.g., the cyan chain) α7 and α8 are unassigned because of their highly flexible structures. The core α2-5 regions in the two protomers are nearly identical, associated with each other in a BH3-in-groove configuration like what we and others previously determined using NMR and crystallography (Czabotar *et al*., 2013; Lv *et al*., 2021).

Each repeating unit has a vortex and two arms (Fig. S12A). The BH3-in-groove core of one asymmetric dimer binds to the core of the other dimer forming the vortex. Each arm contains two α9 helices near the end, one belongs to a protomer in an asymmetric dimer (e.g., the green chain) and the other belongs to another protomer in another asymmetric dimer (e.g., the yellow chain). The interaction between these two α9 helices provides a contact between the two asymmetric dimers in addition to the interaction between the two cores via their adjacent α4 helices (in the green and magenta chains) and the interaction between the two adjacent α6 helices (in the green and yellow chains). The two α9 helices in one arm of the repeating unit (e.g. the green and yellow chains) also interact with the two α9 helices from another repeating unit (e.g., the gray and wheat chains), linking the two neighboring units to form a larger oligomer. The interactions within each repeating unit and between neighboring units would generate a line of Bax oligomer that can curve and form a ring via end-to-end α9-α9 linkage. This hypothesis was confirmed by fitting repeating units into cyroEM structures of Bax polygons extracted from membranes with detergent and stabilized by a protein belt.

Resolution of the repeating unit structure is low, 3.2 Å on average. However, our high-resolution structures of the BH3-in-groove core dimer (blue and marine chains) and the α9 dimer (red and pink chains) can be superimposed on to the respective structures in the repeating unit thereby validated part of the cryoEM structure (Fig. S12B and C). In the structure of a repeating unit (PDB: 9IXU), two α2-5 dimers in the vortex interact with each other partially via two of their four α4 helices. The other two α4 helices do not interact with each other. Instead, they are close to α9 helices in the arms of the neighboring repeating units. Unlike the strong BH3-in-groove and α9:α9 interactions that in fact exist between mitochondria-bound Bax proteins and contribute to Bax oligomeric pore assembly (Bleicken *et al*., 2014; Czabotar *et al*., 2013; Dewson *et al*., 2012; Zhang *et al*., 2016), the contribution of the newly discovered, but relatively mild, α4:α4 and α4:α9 interactions (Zhang *et al*., 2025) to Bax oligomerization in and perforation of the mitochondrial membrane requires further experimental assessment.

The next newly revealed interface in the right arm of the repeating unit is formed between two α6 helices in the cyan and magenta chains shown in Figure S12A, one crossed over from the core dimer on the left side and the other extending from the dimer on the right side. The same interface also exits on the left arm of the repeating unit and is formed similarly between the remaining two α6 helices in the yellow and green chains. Our previous photocrosslinking data with a photoreactive probe attached to Bax residue 134 or 145 can be interpreted by the α6:α6 interface structure (Zhang *et al*., 2010). However, the residues on the other Bax protein that the probes reacted with, and hence in proximity of the probe-attached residues, were not known. According to a disulfide-crosslinking study by Dewson et al, residue 136, 139, 142 or 146 in one Bax molecule is close to the same residue in the other Bax molecule such that each of the residues, when replaced by a Cys, forms disulfide-linked homodimer (Dewson *et al*., 2012). The α6:α6 interface structure in the repeating unit does not fit with these crosslinking data because the Cys residues at each of these locations are too far away to be linked by a disulfide. Similarly, although some α6 helices are close to other α6 helices in our Bax oligomeric pore models from MD simulations (Table S3), their interface structures are different from that in the cryoEM structure. Like the cryoEM interface structure, the simulated interface structure fits with the photo- but not the disulfide-crosslinking data.

The two four-α9 helical bundles, one in each of the two arms of a repeating unit, are perpendicular. If this topographic arrangement occurs in Bax pores in the mitochondrial membrane and one of the helical bundles is spans the membrane and parallel to the membrane normal axis, the other helical bundle must embed in the membrane parallel to the in-plane axis. Otherwise, the membrane must be twisted to accommodate both helical bundles in the same membrane topography, either parallel to the membrane normal or in-plane axis. Most likely all the α9 helices initially insert into the membrane with the same topography, and then a half of them must change their topography if they do not twist the membrane. Alteration of α9 topography after the initial insertion into the membrane was discussed after an EPR-based membrane-bound Bax dimer model proposed opposite topographies for the two α9 helices (Andrews, 2014; Bleicken *et al*., 2014).

There is a large hole in the repeating units encircled by the nonpolar surfaces of the two α2-5 dimers, the two α9 helical bundles, and the four α6 helices. While structures of multiple repeating units can be docked into the EM density maps of Bax polygons, the density maps do not include any lipids, detergents or belt proteins. Hence, how these pore-like structures engage membranes can only be speculated. Our core dimer structure, in contrast, was determined in the presence of lipid/detergent bicelles so it shows the protein-lipid interaction leading to a model in which the core dimer uses its nonpolar surface to cover the nonpolar core of lipid bilayer forming part of the pore wall. Polar residues on the edge of the nonpolar surface bind to the polar lipid headgroups, thereby dictating the orientation of the core dimer relative to the lipid bilayer (Lv *et al*., 2021). On the other hand, our α9 dimer structure determined in lysolipid micelles can be superimposed on two of the four α9 helices in the helical bundle structure revealed by cryoEM and hence is consistent with this α9:α9 interface structure that links adjacent repeating units (Fig. S12C). The other α9:α9 interfaces in the four-helical bundle have different structures than our α9 dimer structure. However, some of our previous and current crosslinking data and MD simulations suggested that the α9 dimer structure is dynamic and could assume other conformations in membranes (Fig. S7B) (Liao *et al*., 2016; Zhang *et al*., 2016). Moreover, inaccessibility of the α9 in the mitochondrial Bax protein to a membrane-impermeable labeling agent and MD simulations of α9 dimers in the model mitochondria lipid bilayer suggest that all α9 dimers assume the same transmembrane topology (Fig. 6A and S6B) (Liao *et al*., 2016; Zhang *et al*., 2016).

The resolution of Bax polygons (oligomeric pores) is even lower (∼5 Å) although some of the secondary structural elements of Bax such as the a2-5 and a9 dimers can be docked into the EM map generating composite models of Bax oligomers. Using Bax tetragon as an example, two adjacent a2-5 dimers can be docked into each of its vertices and a four-a9 helical bundle between two neighboring vertices. Thus, a tetragon contains 16 Bax protomers, same as the number of Bax molecules in our pore models generated by MD simulations. The diameter of the tetragon is ∼83 Å, slightly larger than the diameter of our largest pore (∼70 Å). One reason for the larger diameter and hence circumference of the pore revealed by the EM is that the four-a9 helical bundles are included in the pore wall. In contrast, a9 helices are not located in the pore wall in our pore models. It is worth noting that a large portion of the EM map cannot be docked with any of the remaining Bax regions. This portion may contain detergents that extracted Bax oligomers from cells, lipids that bind to the oligomers, and membrane scaffold proteins that stabilize the oligomers for the EM imaging. As expected, the alignment of the cryoEM structure of a repeating unit and the structure of two adjacent dimers in a pore model from our MD simulations is poor (Fig. 12D).

In summary, our extensive biochemical and biophysical data from Bax proteins in membranes, high-resolution structures of pore components and computational membrane pore models not only are consistent with part of the cryoEM Bax pore structure but suggest more molecular details about the Bax pore assembly in the mitochondrial membrane, thereby enabling future mechanistic structure-function investigation and therapeutics exploration.

(v) *In addition to the contribution of a stable α9 dimer structure to a functional oligomeric Bax pore assembly in the mitochondrial membrane that we have established through this study, the α9 dimer may also contribute to other aspects of Bax life*.

### Dimerization of the α9 helices may occur between soluble Bax proteins

The first structure of a soluble Bax protein determined by NMR shows that the α9 is bound to the BH3-binding groove such that the nonpolar residues that could mediate the α9 dimerization are buried in the groove thereby not available for the dimerization (Suzuki *et al*., 2000). A biophysical study using electron spin resonance (ESR) and double electron-electron resonance (DEER) reveals a fraction of soluble Bax proteins in a conformation that is different from the conformational ensemble determined by NMR (Tsai *et al*, 2015). In particular, the α9 swings out of the BH3 binding groove and is more exposed to solvent. However, other biophysical studies using NMR and/or DEER does not detect this conformation of Bax in solution (Barnes *et al*, 2017; Bleicken *et al*., 2014). The discrepancy could be caused by different experimental protocols used in these studies, e.g., protein purification using different buffers and chromatography methods, and DEER measurement using proteins with different spin-labeling sites. Interestingly, a FRET study of Bax proteins labeled with both donor and acceptor at different sites and microinjected into cells suggests that the α9 is farther away from the α2, a part of the BH3-binding groove before staurosporine was added to induce Bax translocation to the mitochondria (Gahl *et al*, 2014). Thus, Bax protein in solution and in the cytosol can assume a conformation with the α9 out of the BH3-binding groove. If the solvent-exposed α9 interacts with another α9 forming a dimer structure as we determined here, many of the nonpolar residues would be buried in the dimer interface thereby reducing the energy cost of exposing the amphipathic α9 to an aqueous milieu. The resulting α9 dimer would engage the mitochondrial membrane peripherally unless a BH3-only protein interacts with the Bax changing its conformation to favor integration into the membrane (Edlich *et al*, 2011; Gavathiotis *et al*., 2010; Lovell *et al*, 2008). The peripherally interaction with the membrane surface may be mediated in part by the polar residues on the α9 dimer surface that can form hydrogen bonds or salt bridges with the polar lipid headgroups. A previous study showed that the T182I mutation in α9 shortens the residence time of Bax at the mitochondria in healthy cells and hence increases the threshold of apoptotic induction in stressed cells (Kuwana *et al*., 2020). Since the mutation precludes T182 from forming a bifurcated hydrogen bond with T186, it would reduce the stability of α9 and hence the dimerization of α9 and the interaction of Bax with the mitochondria that may be in part mediated by the α9 dimer. Even if the mitochondrial association is mediated by the α9 monomer bound to the membrane peripherally as previously argued (Bleicken *et al*., 2018; Kyrychenko *et al*., 2022), the mutation to isoleucine would eliminate the side chain hydroxyl of Thr^182^ that could otherwise form hydrogen bond with lipid headgroups thereby weaken the a9-membrane interaction.

Bax dimerization and oligomerization can be induced by Bim BH3 peptide in solution as detected by size exclusion chromatography and intermolecular DEER (Gavathiotis *et al*., 2010; Sung *et al*, 2015). Models of Bax dimer and oligomer were created to fit the DEER data, in which α9 helices form an antiparallel dimer instead of the parallel dimer we determined here. Bax dimerization also occurs after interaction with Bim protein in solution or cell lysate as we detected by intermolecular FRET and crosslink, respectively (Chi *et al*, 2020). Whether these soluble Bax dimers contain an interface between the α9 helices and the orientation of the helices in the dimer have not been determined. Based on sequence alignment with the BH3 domain of Bim, the C-terminal sequence of Bim may function as another BH3 domain. The crosslink data suggest that the BH3 domain and the CTS can interact with the BH3-binding groove of Bax protein in cell lysate. Thus, in the Bim-Bax complex it is safe to assume that the α9 helix has been displaced by either the BH3 domain or the CTS of Bim from the BH3-binding groove of Bax thereby available for interaction with another α9 helix in another Bim-Bax complex in the cell lysate.

### Hydrophobicity of α9 dictates insertion of α9 into the MOM that in turn dictates the number of Bax proteins anchored in and perforating the MOM (Chio et al, 2017)

Physicochemical properties of tail-anchors direct tail-anchored proteins to distinct organelles in a cell. A tail-anchor (TA) includes a transmembrane domain (TMD) and a C-terminal element (CTE). A TA with a weakly hydrophobic TMD followed by a weakly basic CTE targets the protein to mitochondria. In contrast, A TA with a moderately to strongly hydrophobic TMD and a weakly basic CTE targets the protein to endoplasmic reticulum, while a TA with a moderately hydrophobic TMD and a strongly basic CTE targets the protein to either mitochondria or peroxisome or both. The hydrophobicity of Bax TMD (α9) is the lowest among the tail-anchored Bcl-2 family proteins that have a TMD. To compare, the grand average hydrophobicity (GRAVY) of the TMD of each of these Bcl-2 family proteins was calculated with ProtParam tool (https://web.expasy.org/protparam/). The TMD’s GRAVY of Bax is 1.384, lower than that of Bcl-2 (1.613), Bcl-W (1.670), Bcl-XL (1.684), Bok (1.800), Bnip3 (1.874), Bak (1.991), Mcl-1 (2.168), and Bik (2.539). The lowest hydrophobicity of Bax TMD may ensure most, if not all, Bax proteins are targeted to mitochondria but not to endoplasmic reticulum or peroxisome once they are activated. The lowest hydrophobicity may also ensure that the inactive Bax proteins remains soluble in the cytosol of healthy cells and not inserted into any membranes. Consistent with this notion increasing the hydrophobicity of the TMD by a mutation such as S184V or S184A resulted in constitutive association of Bax with the mitochondrial membrane, whereas decreasing the hydrophobicity by mutation such as S184K, S184E or S184D abolished Bax translocation from the cytosol to the mitochondria (Nechushtan-Youle 1999). The charge of Bax CTE with residues KKMG, estimated at pH 7.0 with Protein Calculator (https://protcalc.sourceforge.net), is +1.9, which is moderate when compared to Bok (-0.1), Bcl-W, Mcl-1 and Bik (+0.9), Bcl-2 (+1.1), Bcl-XL and Bnip3 (+1.9), and Bak (+2.9). Thus, unlike the hydrophobicity of TMD, the charge of CTE seems less important in specifying the localization of these proteins.

Bax α9 when fused with GFP localized the fusion protein to mitochondria (Ref 57, 58 of Manon 2022 Biomolecules review; the following ref’s are #’ed similarly). Bax with the α9 however did not localize to mitochondria (59). Replacing the Bax α9 with the Bcl-XL α9 localized the chimeric protein to mitochondria, while replacing the Bcl-XL α9 with the Bax α9 prevented the chimera to localized to mitochondria. This study shows that the Bcl-XL α9 contains all the information required for mitochondrial localization whereas Bax a9 does not. The charged “X-domain” before and the positive charged residues after the TMD (Manon 2022 Biomolecules review) are important for targeting Bax and other Bcl-2 proteins to the mitochondria. While Bax lacks the X-domain, Bcl-2 and -XL have. However, the data above also can be explained by the fact that Bax α9 can fold into the BH3 binding groove of Bax and the possibility that Bax α9 may also fold into the groove of Bcl-XL. In contrast, the Bcl-XL α9 may not fold into the Bcl-XL groove in the full-length Bcl-XL protein and the groove of Bax in the chimeric protein. Consistent with this explanation, deleting or changing Ser^184^ in Bax α9 to Ala or Val induced a constitutive localization of Bax to mitochondria (49) as the mutations would loosen the interaction of α9 with the groove since they remove a hydrogen bond with Asp^98^ in the groove or generate steric clashes or lose van de Waals contacts with the groove. There is Pro^168^ in the short loop between α8 and α9 that controls whether the α9 is oriented into the groove. In the soluble Bax structure, this proline residue is in cis configuration and the α9 is oriented into the groove. Substitution of the proline with alanine that is almost exclusively in the trans configuration increased Bax localization to mitochondria in human glioblastoma cells and released more cytochrome C (71). Intriguingly a peptidyl-prolyl cis-trans isomerase Pin1 could interact with Bax and modulate its mitochondrial localization (76) Whether the integration process can be modulated by the α9 dimerization requires future studies.

(vi) *To interfere Bax pore assembly, which step or interaction is the best point of interference?* (Andrews, Gavathiotis, Moldoveanu, reviews) As an alternative or addition to targeting antiapoptotic proteins, such as the Bcl-2-targeting Venetoclax approved for CLL and AML treatments, targeting Bax could provide new treatments for cancer, cardio and neural diseases and benefit tissue engineering and transplantation (Pogmore et al J Med Chem 2021; Zacharioudakis and Gavathiotis TiBS 2022;). So far, a few small molecule modulators have been developed based on Bax structures in solution and mapping of their binding sites in soluble Bax using NMR and H/D exchange mass spectrometry. Structures of Bax in a lipid environment reveal new binding sites for many more small molecules since extensive biophysical and biochemical studies have shown that membrane bound or embedded Bax proteins have different conformations and surface features than the soluble proteins. This has been demonstrated by that the structure of the a2-5 regions bound to lipid/detergent bicelles is different from that of the same regions in crystals. Now we have the structures of a9 dimer in detergent micelles and Bax oligomers extracted by detergents and associated with belt proteins that reveal potential binding sites for new small molecules. For example, the interface between the lipid bilayer and the a2-5 dimer could be targeted by a small molecule such that the protein wall could be detached from the membrane preventing Bax pore formation. The proof of principle has been provided by the effect of the R89E mutations that inhibits the a2-5 dimer interaction with the lipid bilayer and thereby the Bax pore formation. The a9 dimer interface could be another target. Interestingly the GxxxG or similar motifs for transmembrane helical dimerization was proposed as potential drug targets although peptides but not small molecules were believed to be a viable choice. Here we show that point mutations such as I175A and T182I inhibit the a9 dimerization and the Bax pore formation suggesting that small molecules could be viable alternatives. It could be questioned that the a9 is partially buried in the soluble Bax structures and hence not available for small molecule binding. However, since S184 in the a9 of Bax in cytosol can be accessed by Akt for phosphorylation, the a9 can be partially exposed for small molecules to bind. Also, biophysical studies revealed conformational dynamics for soluble Bax proteins that in part exposed the a9 to the cytosol at least transiently. Critically, the interfaces in these dimers formed by part of Bax in nonnative membrane environments have been extensively verified specially by site-directly crosslinking experiments. It is of a high confidence that they exist in the dimers and oligomers form by the full-length Bax proteins in native mitochondrial membranes. Similarly, the new interfaces revealed by the cryoEM structures of Bax oligomer provide more drug targets if one can prove their relevance to the native Bax pore structures in the mitochondrial membranes. Moreover, recent studies revealed that the size and formation kinetics are different between Bax and Bak pore, the existence of Bax/Bak hybrid pores (Cosentino *et al*., 2022). These features give functional difference between the pores such as they induce apoptosis or inflammation. Since the pore size is controlled by the abundance of active Bax and Bak proteins in the membrane, the rate of their diffusion and productive binding interaction, and the affinity between the pore-forming proteins or the interfaces between the pore-building blocks. Thereby, targeting different interfaces such as the a2-5 or a9 dimer interfaces may provide different ways to control pore size and poration kinetics, which in turn provide different therapeutic outcomes (Andrews, 2022).

## Materials and Methods

### Protein expression and purification

For Bax (α9), *E. coli*-codon optimized DNA encoding residues 166-192 (α9) of human Bax with N-terminal 9×His tag and negative charged E3 tag (EIAALEK, EIAALEK, EIAALEK) was synthesized (GenScript). The Bax (α9) fusion protein was expressed in inclusion bodies. The cells were resuspended and lysed by high-pressure homogenizer in Buffer A (25 mM HEPES, pH 7.5, 150 mM NaCl). After lysis, the inclusion bodies was collected by centrifugation at 40,000 × *g* for 30 min and resuspended in buffer B (25 mM HEPES, pH 7.5, 150 mM NaCl, 2% sarkosyl and 0.1% DPC) by stirring at 4 °C overnight. The supernatant was then subjected to Ni-NTA purification in buffer B, and followed by sequential washing with imidazole gradient (10 mM and 50 mM imidazole, respectively) in Buffer C (25 mM HEPES, pH 7.5, 150 mM NaCl, 0.1% DPC). The protein was eluted using 500 mM imidazole in Buffer C. The sample was concentrated and subjected to size exclusion using the Superdex200 10/300GL column (GE Healthcare) in Buffer C to improve the purity and homogeneity. The final yield of Bax (α9) was about 5 mg from 1 L of cell culture.

### Characterization of Bax (α9) oligomeric states by crosslinking

The oligomeric state of the Bax (α9) in DPC micelles was also examined by chemical crosslinking using BS^3^. The mixtures of 50 μL 100 μM Bax (α9) in DPC micelles with 4 μL 50 mM BS^3^ stock solution were incubated at room temperature for 20 min, and quenched by the addition of 1 M Tris pH 8.0. The reaction mixtures were analysed using SDS–PAGE, and visualized by Coomassie brilliant blue staining.

### NMR data acquisition, processing and analysis

All NMR experiments were conducted on Bruker spectrometers operating at ^1^H frequency of 900 MHz, 600 MHz and equipped with cryogenic probes. NMR spectra were processed using NMRpipe (Delaglio *et al*, 1995) and analyzed using XEASY (Bartels *et al*, 1995) and CcpNmr (Vranken *et al*, 2005). Sequence-specific assignment of backbone amide resonance (^1^H^N^, ^15^N, ^13^Cα, and ^13^C’) was accomplished using a series of gradient-selected, TROSY-enhanced triple resonance experiments, including HNCO, HN(CA)CO, HNCA, HN(CO)CA and HNCACB (Salzmann *et al*, 1999). The NMR data was collected with a uniformly (^15^N, ^13^C, ^2^H)-labeled sample on a 600 MHz spectrometer. In addition, a 3D ^15^N-edited NOESY-TROSY-HSQC experiment (with 150 ms NOE mixing time) was performed to validate the assignment. Protein aliphatic and aromatic side chain resonances were assigned using a combination of a 3D ^15^N-edited NOESY-TROSY-HSQC and a ^13^C-edited NOESY-HSQC recorded on a 900 MHz spectrometer. The proteins used for these experiments were (^15^N, ^13^C)-labeled. Stereospecific assignments of methyl groups of leucine and valine were determined by a ^1^H-^13^C HSQC spectrum recorded on a 15% ^13^C-labeled sample.

To determine the intermolecular distance restraints, a mixed sample in which half of the monomers were ^15^N, ^2^H-labeled and the other half was ^13^C-labeled was used to record a 3D ^15^N-edited NOESY-TROSY-HSQC (τ_NOE_ = 250 ms) to obtain the exclusive NOEs between the ^15^N-attached protons of one monomer and aliphatic protons of the neighboring monomers.

All the experiments for Bax (α9) in DPC micelles were performed at 25 ℃. And the detergents were deuterated for the above mentioned NOESY experiments.

### Structure calculation

Structure was calculated using the program XPLOR-NIH. For Bax (α9), the monomer structure was first derived using intramonomer restraints and backbone dihedral restraints, determined from chemical shifts using the TALOS+ program. We then used the intermonomer contacts from the mixed sample with differentially labeled monomers to derive the dimer structure. A negative control sample was used to confirm the intermonomer NOEs. PRE analysis was performed on a mixed sample containing ∼1:1 ratio of ^15^N-labeled Bax (α9) and ^14^N Bax (α9) tagged with MTSL at positions T172, G179 and W188.

### Structure validation by PRE measurement

To verify the structures calculated above, PRE analysis was performed to obtain intermonomer information independent of NOEs. A cysteine mutation is introduced for labeling the proteins with MTSL ((1-oxyl-2,2,5,5-tetramethyl pyrroline-3-methyl) methanethiosulfonate). Three Cys mutations were introduced separately at positions 172, 179 and 188. The cells of (^15^N) labeled Bax (α9) and the cells of ^14^N Bax (α9) with a Cys mutation were mixed prior to purification. After micelle reconstitution, MTSL was added to the sample at a final ratio (MTSL to Bax (α9)) of 10:1 and the mixture was incubated at 4 ℃ overnight. The sample was passed through a PD-10 desalting column to remove free MTSL before the recording of ^1^H-^15^N TROSY-HSQC spectrum. Sodium ascorbate was added to recover the NMR peak intensity.

### Activity assays for the structure-guided Bax mutants in the mitochondrial membranes

Bax dimerization via a9 was detected using disulfide crosslinking of single-Cys Bax in the isolated Bax-/-/Bak-/- mitochondria as previously described (Lv *et al*., 2021; Zhang *et al*., 2016) as well as Bax-mediated liposome permeabilization. The mitochondrial outer membrane permeabilization by Bax was monitored by following SMAC-mCherry release from the mitochondrial intermembrane space in digitonin-permeabilized BMK Bax-/-/Bak-/- cells as previously described (Lv *et al*., 2021), but additional steps were added to further solubilize the released and unreleased mitochondrial proteins and remove the insoluble materials that scatter light thereby interfering with the mCherry fluorescence measurement. Mainly, 130 μl of the permeabilized cells (1,300,000 cells total) was incubated with 50 nM Bax WT or mutants and/or 5 nM cBid in Eppendorf tubes at 37°C for various times with gentle mixing every 20 min. The tubes were centrifuged at 4,200g and 25°C for 10 min. The supernatant and pellet fractions were collected, and their volume was adjusted to 250 μl with cell buffer (20 mM HEPES-KOH pH 7.4, 250 mM sucrose, 150 mM KCl, 2 mM MgCl_2_, 1 mM EDTA) plus protease inhibitors (0.1 μg/ml Leupeptin, 0.1 μg/ml Pepstatin, 140 μg/ml Aprotinin). Triton X-100 (0.5% v/v) was added to the fractions followed by an incubation on ice for 20 min. The tubes were centrifuge at 4°C and 16,363g for 10 min. The clear supernatants were transferred to cuvettes as 250-μl aliquots and they corresponded to the original supernatant and pellet fractions that contain the SMAC-mCherry proteins that were released by Bax and/or cBid and that were retained in the mitochondria, respectively. The fluorescence intensities of these SMAC-mCherry proteins were measured using ISS PC1 Photon-Counting Spectrometer with red-edge excitation and blue-edge emission to reduce Rayleigh scattering light contamination. Thus, the samples were excited with a xenon arc lamp (I = 18 A) at 545 nm; emission spectra were recorded from 565 to 660 nm with 1-nm increments; slit widths of excitation and emission were 4.8 and 2.0 mm, and with the dispersion of both monochromators = 8 nm/mm the bandpass were 38.4 and 16 nm, respectively; sample temperature was maintained at 25°C; the emission intensity at each wavelength was measured by a photomultiplier tube (PMT) for 2 s during which the PMT collected the emitted photons and converted to an electrical current every 0.1 s, and the resulting 20 electrical currents were integrated to give an emission intensity with a standard error; for each sample, the emission spectrum from each sample was measured twice. The mCherry fluorescence under each spectrum was integrated from 580 to 650 nm to further minimize the contribution from Rayleigh scattering light. SMAC-mCherry release was calculated by dividing the mCherry fluorescence of the supernatant fraction by the total mCherry fluorescence of the supernatant and pellet fractions. The data was normalized by dividing the fraction of SMAC-mCherry release of each sample by the fraction of SMAC-mCherry release of Bax WT and cBid at 60 min.

### Molecular dynamics simulations and modeling of oligomeric Bax pores

#### Model preparation

An oligomeric pore model was constructed from sixteen truncated Bax molecules (residues 48-192), dimerizing via the α2-5 regions and the α9 regions, respectively. The dimeric conformations were constructed from the NMR structures of α2-5 dimer and α9 dimer (PDB ID: 6L8V and 6L95, respectively) using Maestro (Schrödinger, LLC). The initial configuration is displayed in Fig S6A, where the α2-5 dimers form the pore wall, the α9 dimers insert into the membrane, and they are connected by the α6-8 regions.

We built the protein-membrane complex system using the membrane builder of CHARMM-GUI (Jo *et al*, 2008), and removed lipids within the pore. The Bax oligomer was embedded in a lipid bilayer consisting of 46.5, 28.4, 8.9, 8.9 and 7.3% of L-α-phosphatidylcholine (PC), L-α-phosphatidylethanolamine (PE), L-α-phosphatidylinositol (PI), L-α-phosphatidylserine (PS) and cardiolipin (CL) in mole fraction, respectively. The aqueous phase contains 132,818 TIP3P water molecules, 0.06 M KCl as counter ions, together with the protein-membrane complex totaling near 631,822 atoms in a periodic box 259×259×103 Å^3^.

#### Simulation Setup

All simulations were performed with the CHARMM36m-cmap force field (Best *et al*, 2012). Each system went through energy minimization, multi-step isothermal-isovolumetric (NVT) equilibration scheme of total 1,200 ps with the force constant of dihedral restraint on proteins and lipids gradually reduced from 250 kcal/mol to 0 kcal/mol in six steps. Then, the system continued to relax in microsecond MD simulations under isothermal-isobaric ensemble with semi-isotropic pressure coupling (NPγT) using the AMBER16 package (D.A. Case *et al*, 2018) with GPU acceleration. The NPγT ensemble was set with 310 Kelvin, 1 bar, Langevin dynamics thermostat and semi-isotropic Monte Carlo barostat and a time step of 2 fs. All lengths of bonds to hydrogen atoms in protein or lipid molecules were constrained with SHAKE. The particle mesh Ewald (PME) technique was used for the electrostatic calculations. The van der Waals and short-range electrostatics were cut off at 12.0 Å with switch at 10.0 Å. Each system was simulated for over 700 ns. Each simulation was replicated ten times.

#### Data analysis

Data from ten simulation replicas were analyzed with build-in analysis tool, “cpptraj”, (Roe & Cheatham, 2013) in AMBER package, then further processed and plotted using matplotlib (Hunter, 2007). Structures are shown with PyMOL (Schrödinger, LLC). Protein-protein contacts are counted by residues whose Cα-Cα distance is within 6 Å. Contacts between negatively charged phospholipids and protein residues are counted by their P-Cα distance within 10 Å.

## Data availability

NMR structure coordinates have been deposited in Protein Data Bank under accession number 6L95 (https://www.rcsb.org/structure/6L95). ^1^H, ^13^C, and ^15^N chemical shifts have been deposited in Biological Magnetic Resonance Bank under accession number yyyyy (https://bmrb.io/data_library/summary/index.php?bmrbId=yyyyy). Oligomeric pore structural models of 16 Bax a2-9 regions in a mitochondria-mimic lipid bilayer have been deposited in Protein Data Bank under accession number xxxx (https://www.rcsb.org/structure/xxxx)

## Acknowledgements

We thank the staffs from Large-scale Protein Preparation, Nuclear Magnetic Resonance and Mass Spectrometry Systems at National Facility for Protein Science in Shanghai, Zhangjiang Laboratory, China for technical support and assistance in data collection and analysis. This work was supported by grants from National Key R&D Program of China (2017YFA0504804), Key Research Program of Frontier Sciences, CAS (QYZDB-SSW-SMC043) to B. O., by grants from US National Institutes of Health (R01GM062964), OCAST (HR16-026) and Presbyterian Health Foundation to J. L., by an Institutional Development Award from the National Institute of General Medical Sciences of US National Institutes of Health (P20GM103640), and by a foundation grant from the Canadian Institutes of Health Research (FDN 143312) to D. W. A. D. W. A. holds the Tier 1 Canada Research Chair in Membrane Biogenesis.

## Author contributions

J. L., B. O., and C. L. conceived and directed the project; B. O. designed NMR experiments; B. O. and J. L. designed mutations and functional assays; L. Z., F. L. and L. D. prepared Bax (α9) proteins for NMR, collected and analyzed NMR data and determined the Bax (α9) structure with the help from B. O.; L. Z., F. L. and L. D. analyzed PRE data and built the structural model for micelle-embedded Bax (α9); F. Q. and J. D. R. prepared Bax proteins, performed liposomal dye release experiments and analyzed the data with J. L.; Z. Z. generated Bax mutant plasmids, performed crosslinking experiments, liposomal dye and mitochondrial protein release experiments, and analyzed the data with J. L.; J. P. purified mouse liver mitochondria; C. L. performed MD simulations of Bax oligomers in lipid bilayer; J. L., B. O., C. L. and D. W. A. interpreted the data and wrote the paper; All authors edited the article.

---

**Figure S1.**
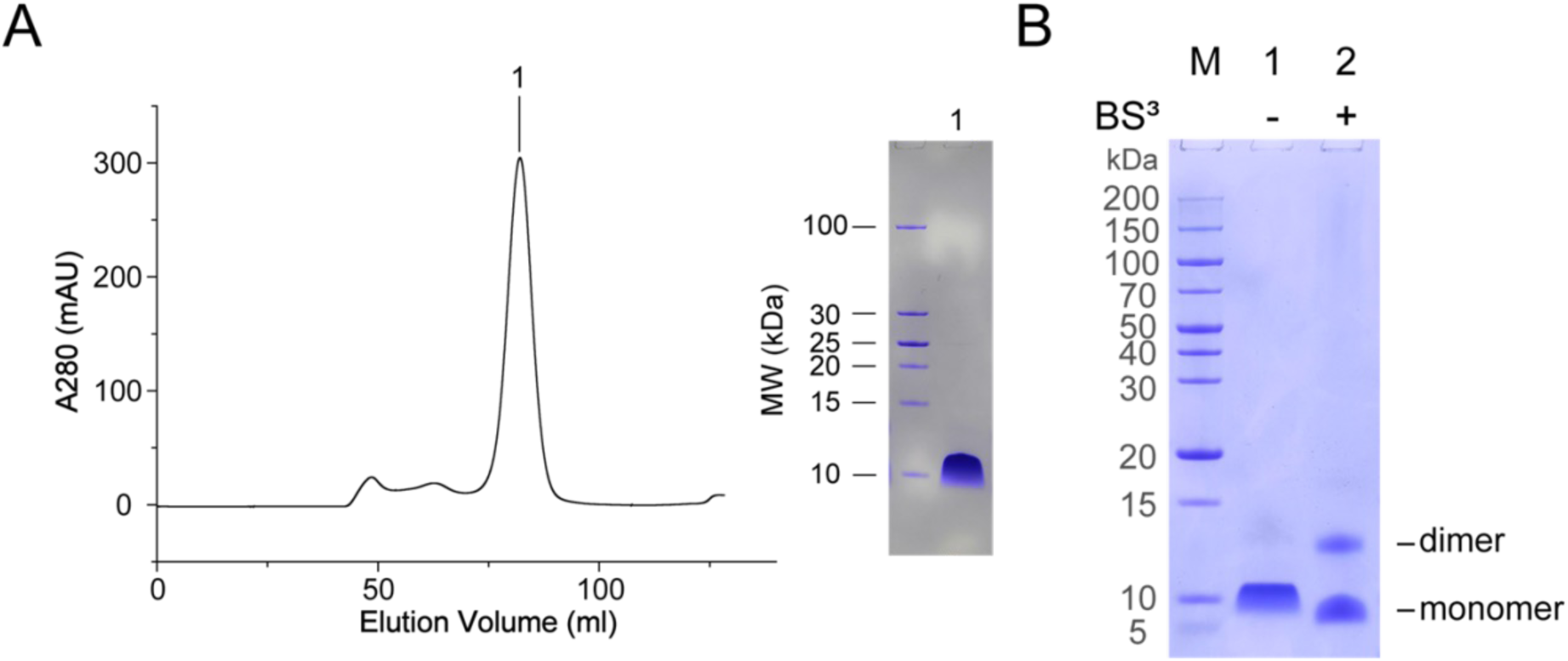
Purification and Crosslinking of Bax (α9) **A**. Size exclusion chromatography of Bax (α9) in DPC micelles (left). The product was verified by SDS-PAGE (right). **B.** SDS-PAGE analysis of the BS3 crosslinking of Bax (α9) NMR sample.

**Figure S2.**
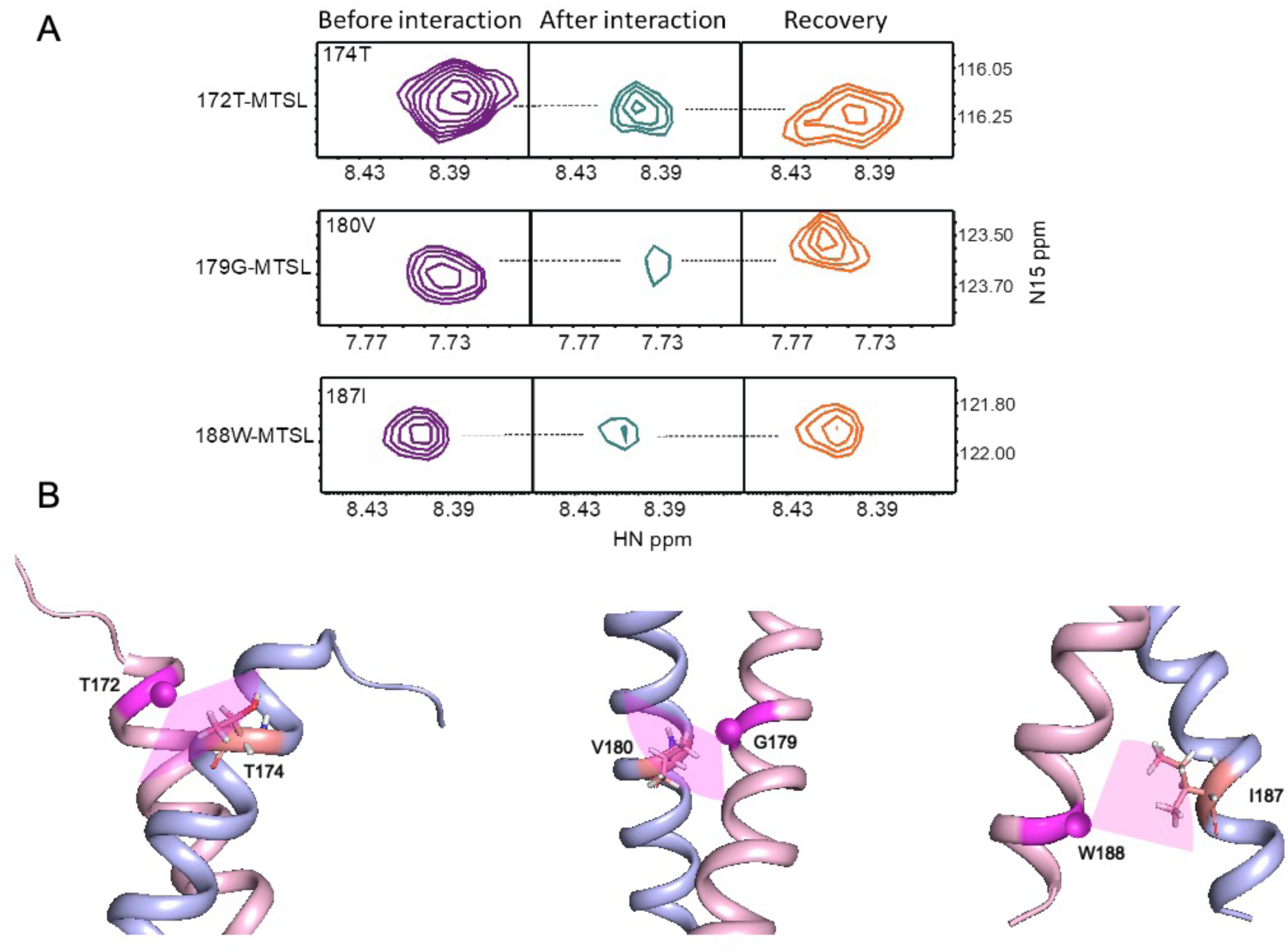
Structure verification of Bax (α9) dimer using paramagnetic probes. **A.** Specific broadening of 1H-15N correlation peaks by paramagnetic probe MTSL and recovery of the broadened peaks upon addition of ascorbic acid. **B.** Interchain PRE analysis of Bax α9 dimer in micelles. Mapping the PREs onto the structure of Bax α9 dimer in bicelles, showing the backbone amide protons (residue represented by a salmon stick in each structure) in chain A (light blue) with strong PREs to the nearby spin probe (represented by a magenta sphere in each structure) in chain B (pink). The sample consisted of an ∼1:1 mixture of [15N]-labeled Bax α9 and [14N]-labeled Bax α9 with a MTSL spin probe at T172C (left), G179C (middle) or W188C (right).

**Figure S3.**
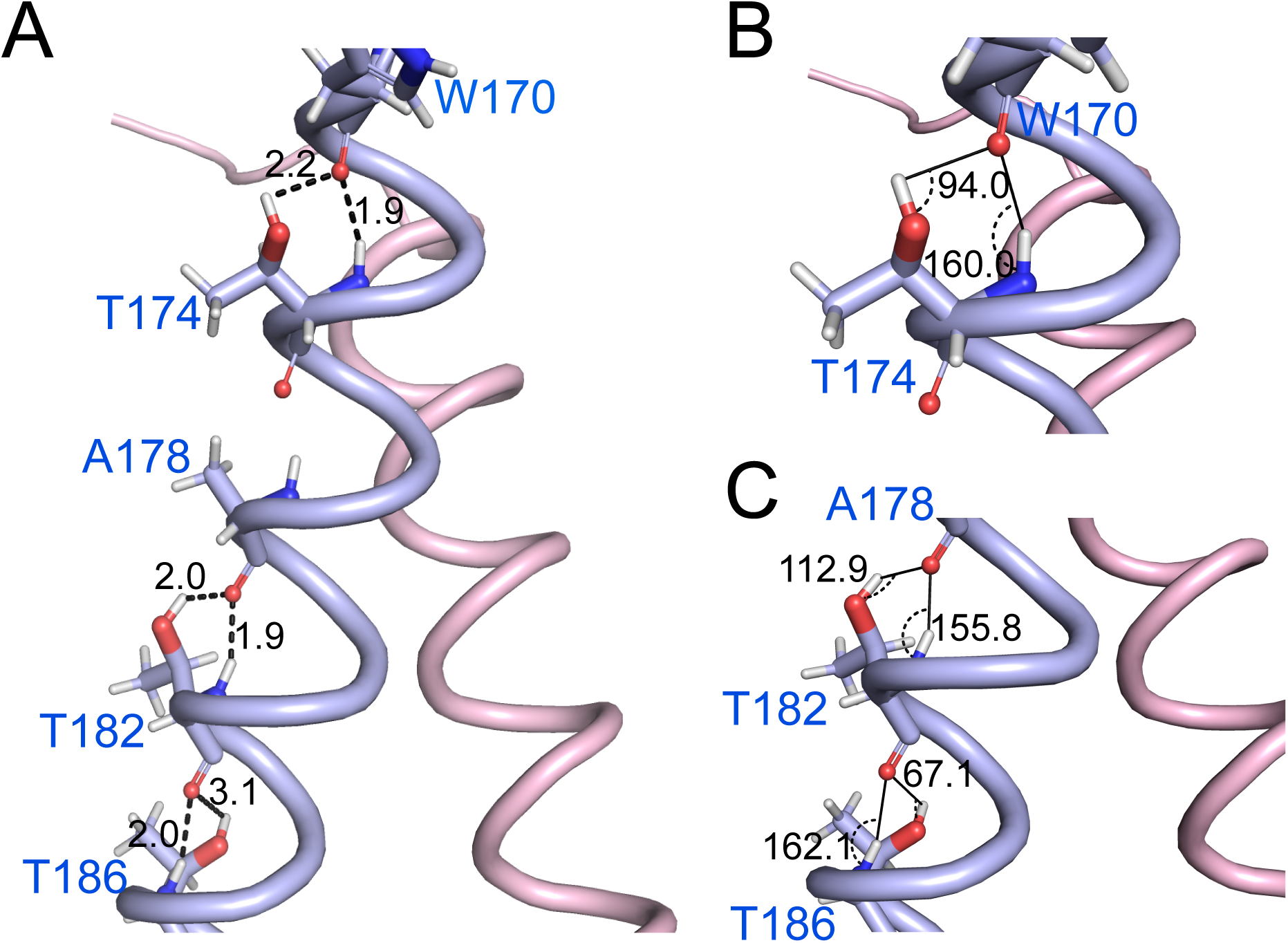
Bifurcated hydrogen-bonds in α9 dimer. **A**. The residues involved in the bifurcated H-bonding are shown as sticks. The dashed lines depict the mainchain-sidechain and mainchain-mainchain H-bonds, each with the donor-acceptor distance between the donor hydrogen atom and the acceptor hydroxyl or carbonyl oxygen atom indicated in angstroms. **B.** The donor-acceptor-acceptor antecedent angle of each H-bond is indicated in degrees.

**Figure S4.**
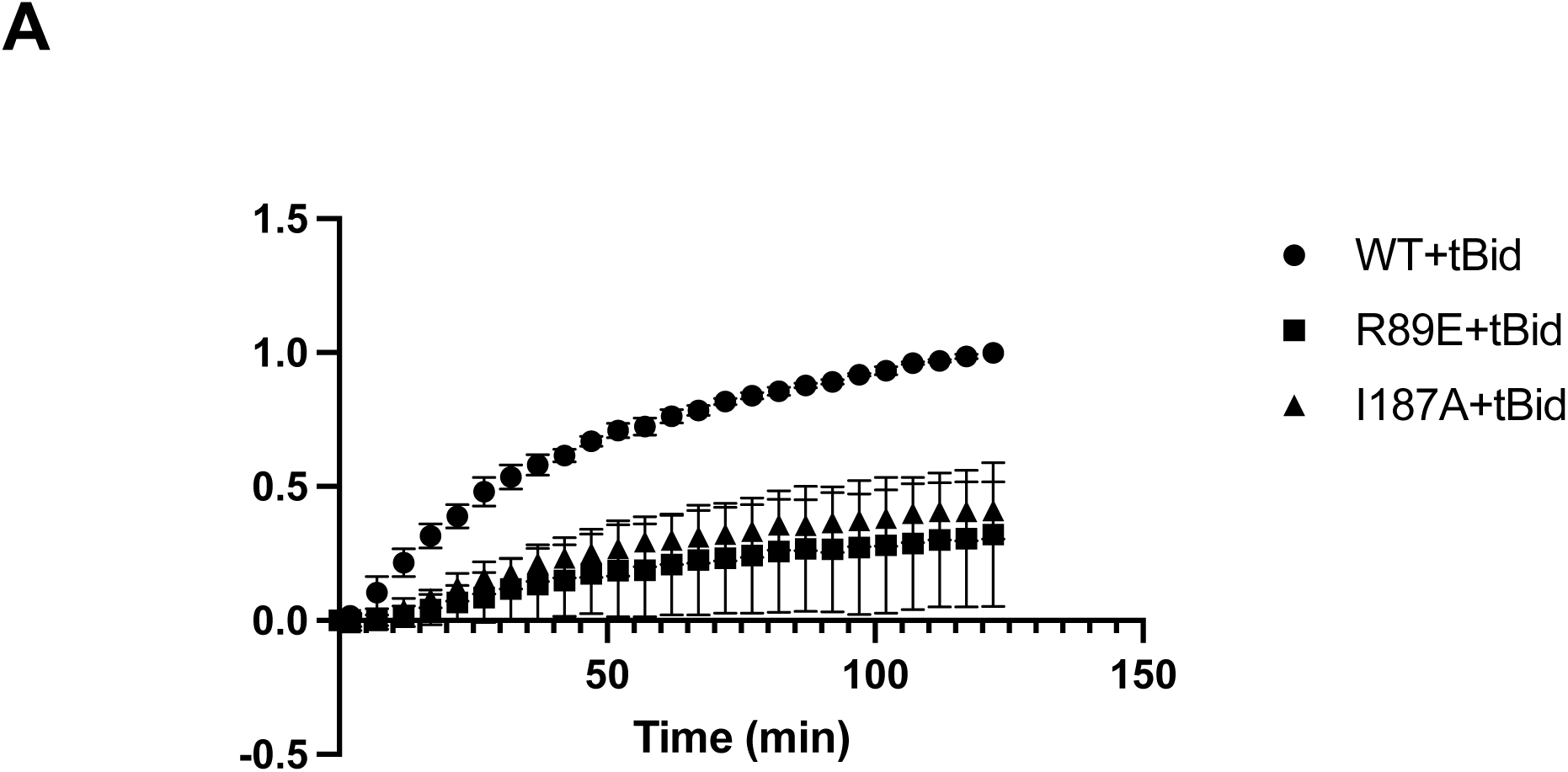
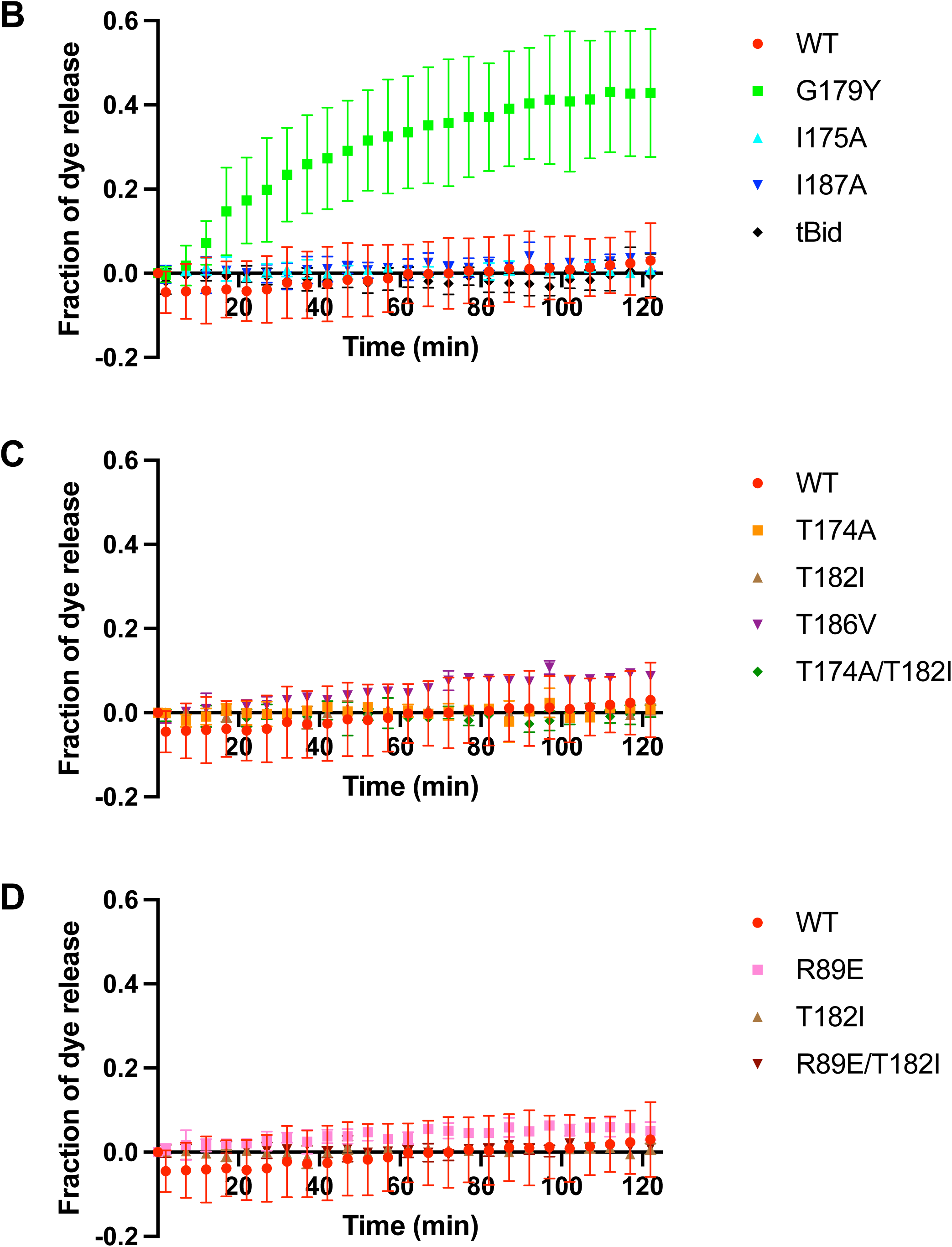
Liposomal dye release by a lower concentration of Bax, and by Bax or tBid alone. (**B**) Fluorescent dye release from the liposomes by 25 nM wild-type (WT) or mutant Bax in the presence of 15 nM tBid was measured. The data obtained from n = 3 independent replicates were processed and presented as that in Figure 4. (**B-D**) Fluorescent dye release from the liposomes by 100 nM wild-type (WT) or mutant Bax, or 15 nM tBid was measured. The data obtained from n = 2 to 5 independent replicates were processed and presented as that in Figure 4.

**Figure S5.**
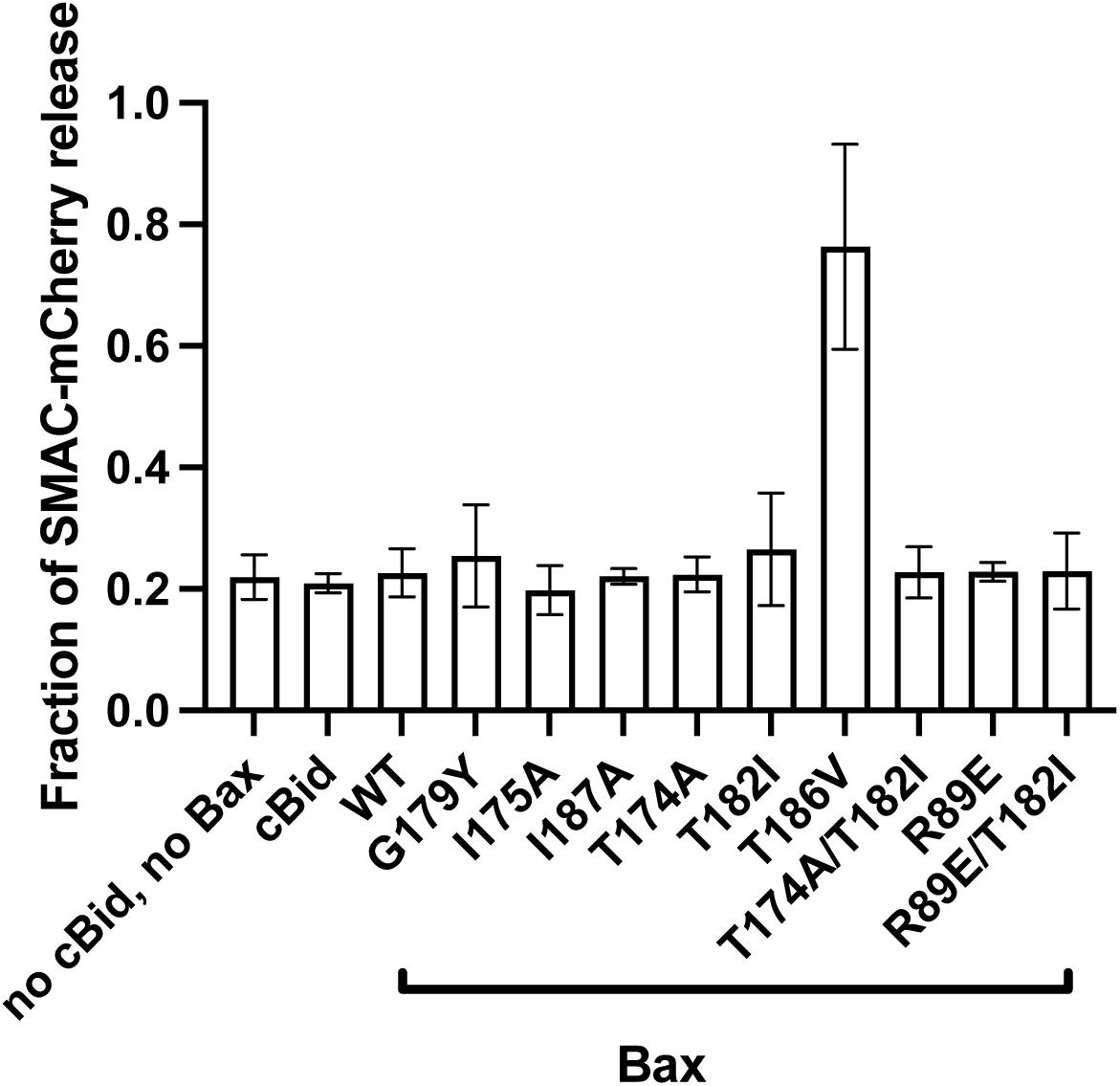
Mitochondrial SMAC-mCherry release by no proteins, and single Bax wild-type or mutant proteins or tBid. The SMAC-mCherry release by no proteins or individual proteins by the end of a 60-min incubation was measured as described in Figure 5. The data from n = 3 or more independent replicates were processed and shown as the means and with the error bar representing standard error of the mean.

**Figure S6.**
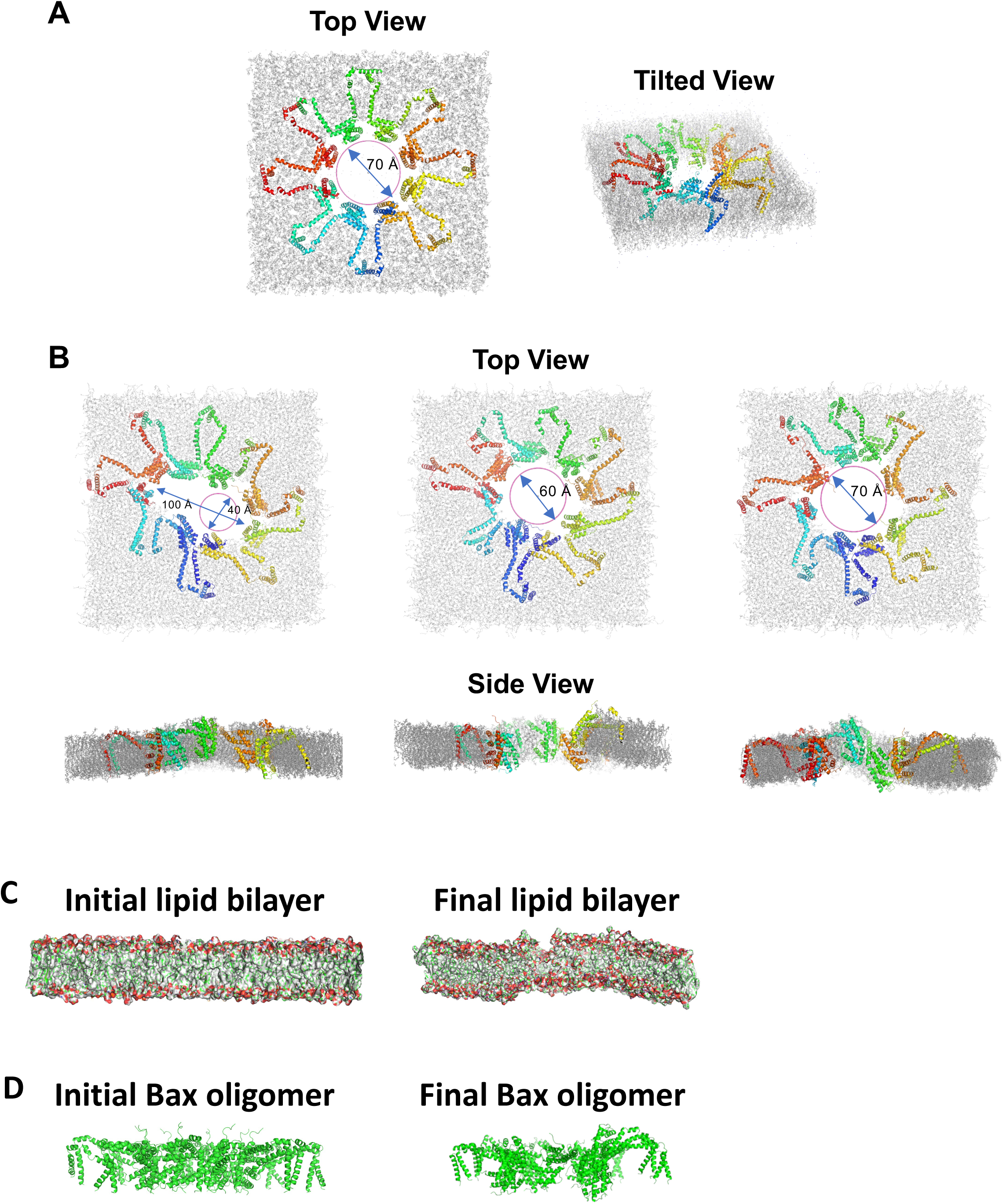
Bax pores of various sizes and associated membrane curvature observed by the end of MD simulations. Top and tilted views of the initial pore structure for MD simulations. **A.** Representative final pore structures from MD simulations. The top views show the elliptical (left) and circular (middle and right) pores evolved from the initial pore by the end of ten independent 700-ns MD simulations. The major and minor axes of ellipse and the diameter of circle are indicated in Å. The side views show membrane curvature around the pores. **B.** The lipid bilayers associated with the initial and final pore structures show evolution of membrane curvature during MD simulations. **C.** The initial and final structures of the Bax oligomer show alteration of conformation during MD simulations.

**Figure S7.**
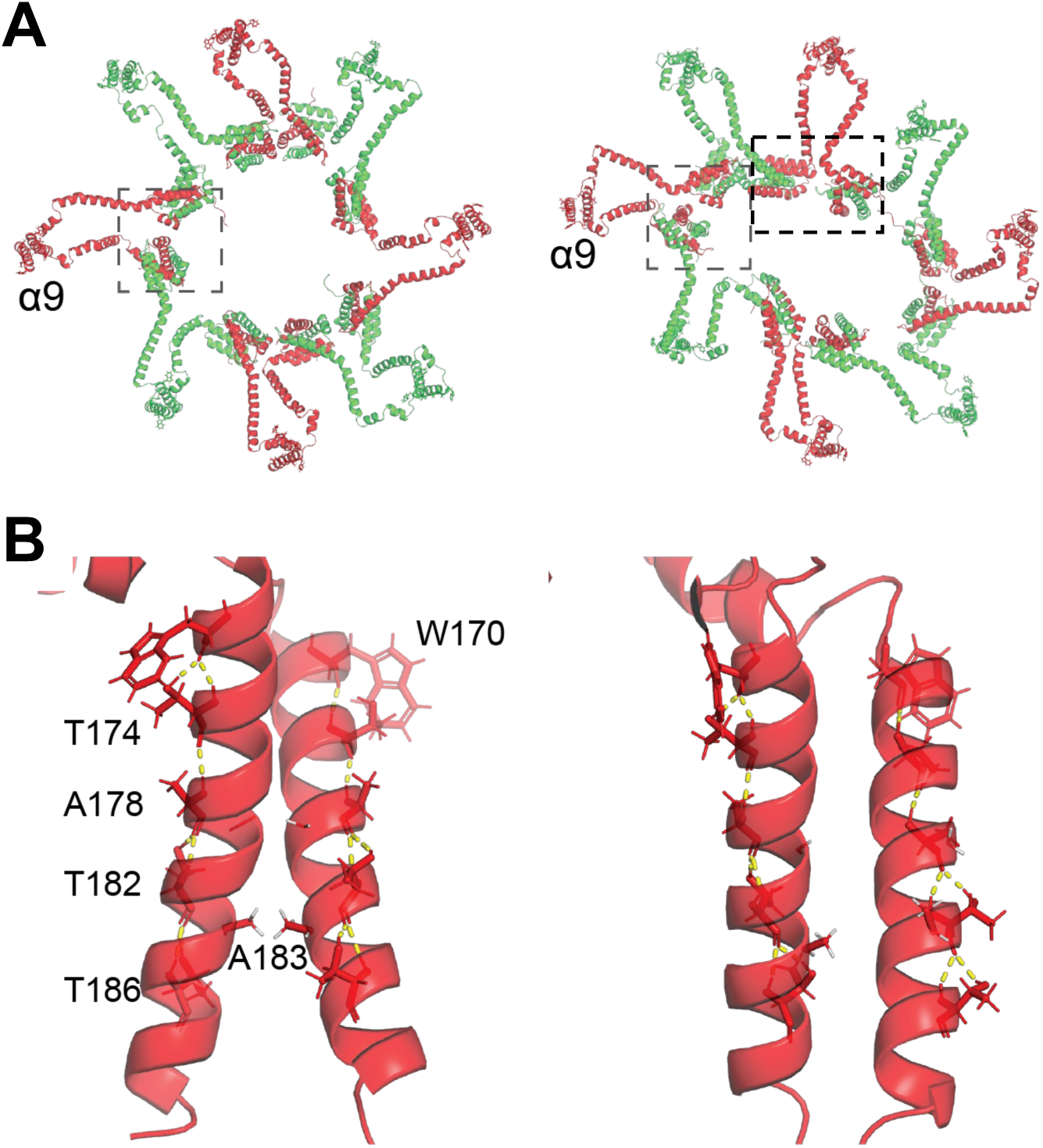
Various configurations of pore components generated by the end of MD simulations. **A.** Circular (left) and elliptical (right) pores. Representatives of adjacent α2-5 dimers are enclosed in rectangles. Representatives of α9 dimer are indicated by ‘α9’. **B.** Configurations of α9 dimer indicated in (A) and bifurcated H-bonds in α9.

**Figure S8.**
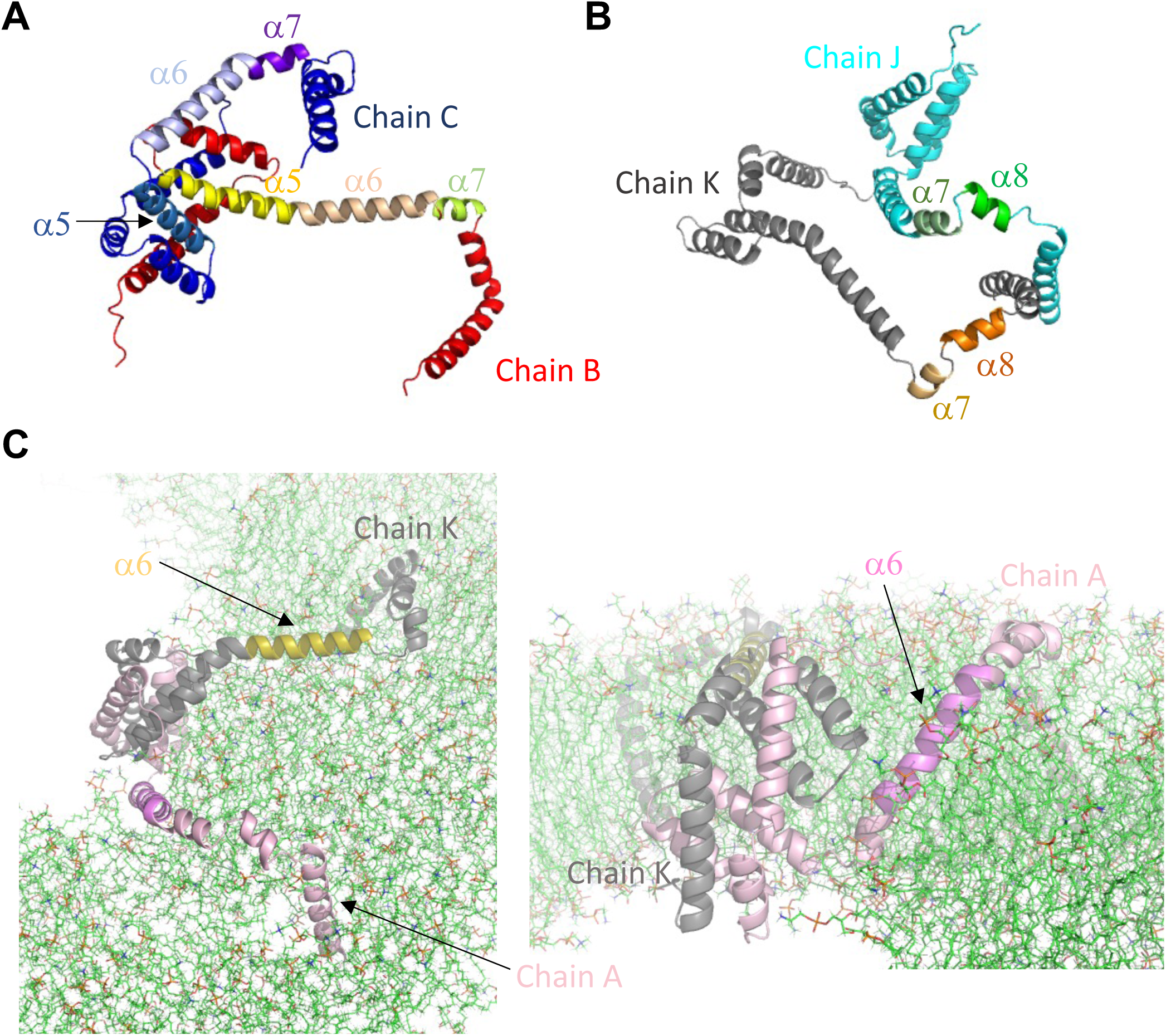
A variety of conformations and membrane topographies of α5-8 regions observed by the end of MD simulations. **A.** Typical conformations of α5-7 regions. **B.** Typical conformations of α7-8 regions. **C.** Typical membrane topographies of α6.

**Figure S9.**
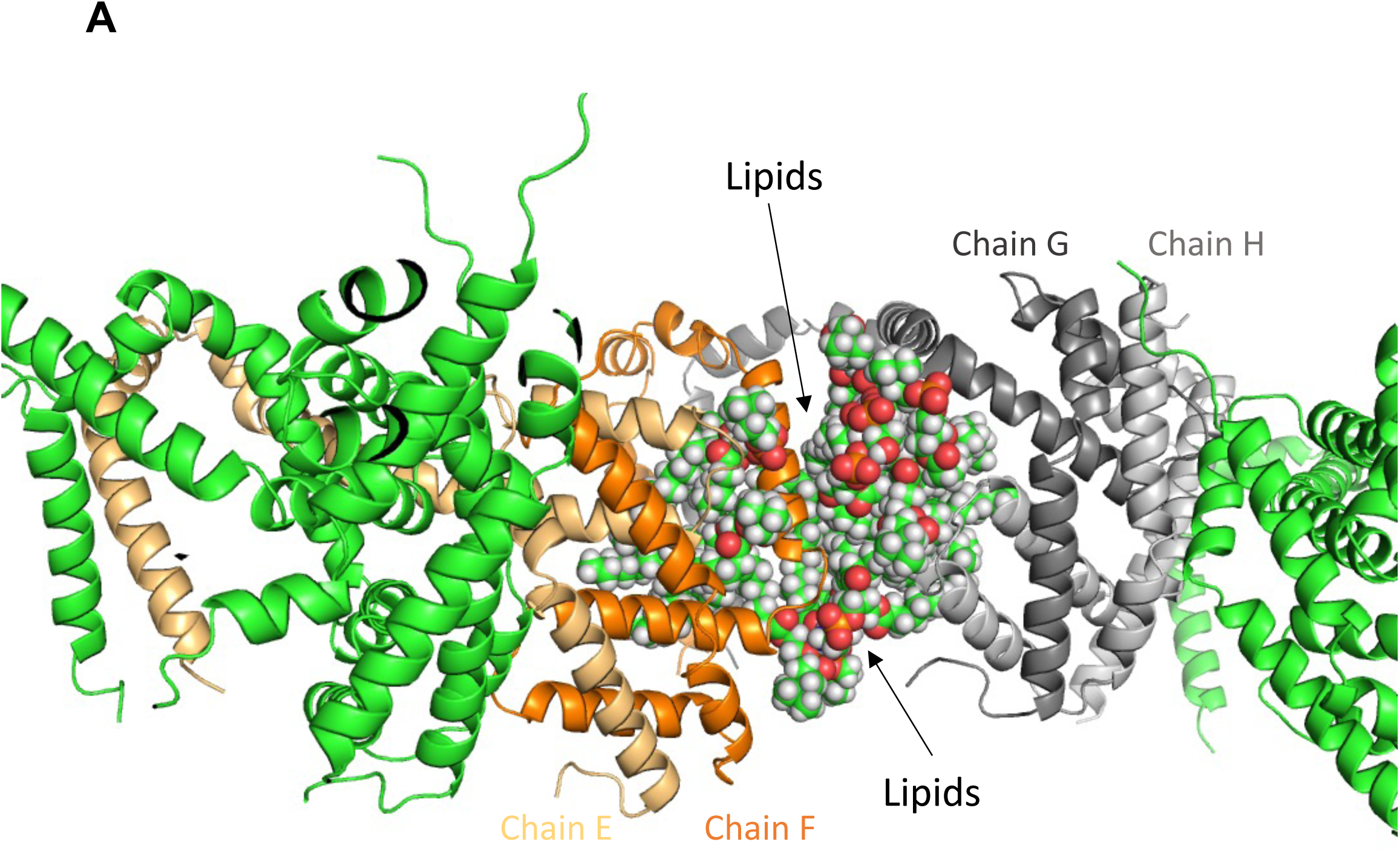
Lipids that fill a gap between adjacent a2-5 dimers in the pore wall observed by the end of MD simulations. POPC, POPC, POPC, POCL, POPC, POPE, POPC, POPC, POPC, and POPC that fill a typical gap in the protein pore wall are shown as spheres with the polar or charged headgroups in red, and the nonpolar acyl chains in green (for carbons) and white (for hydrogens).

**Figure S10.**
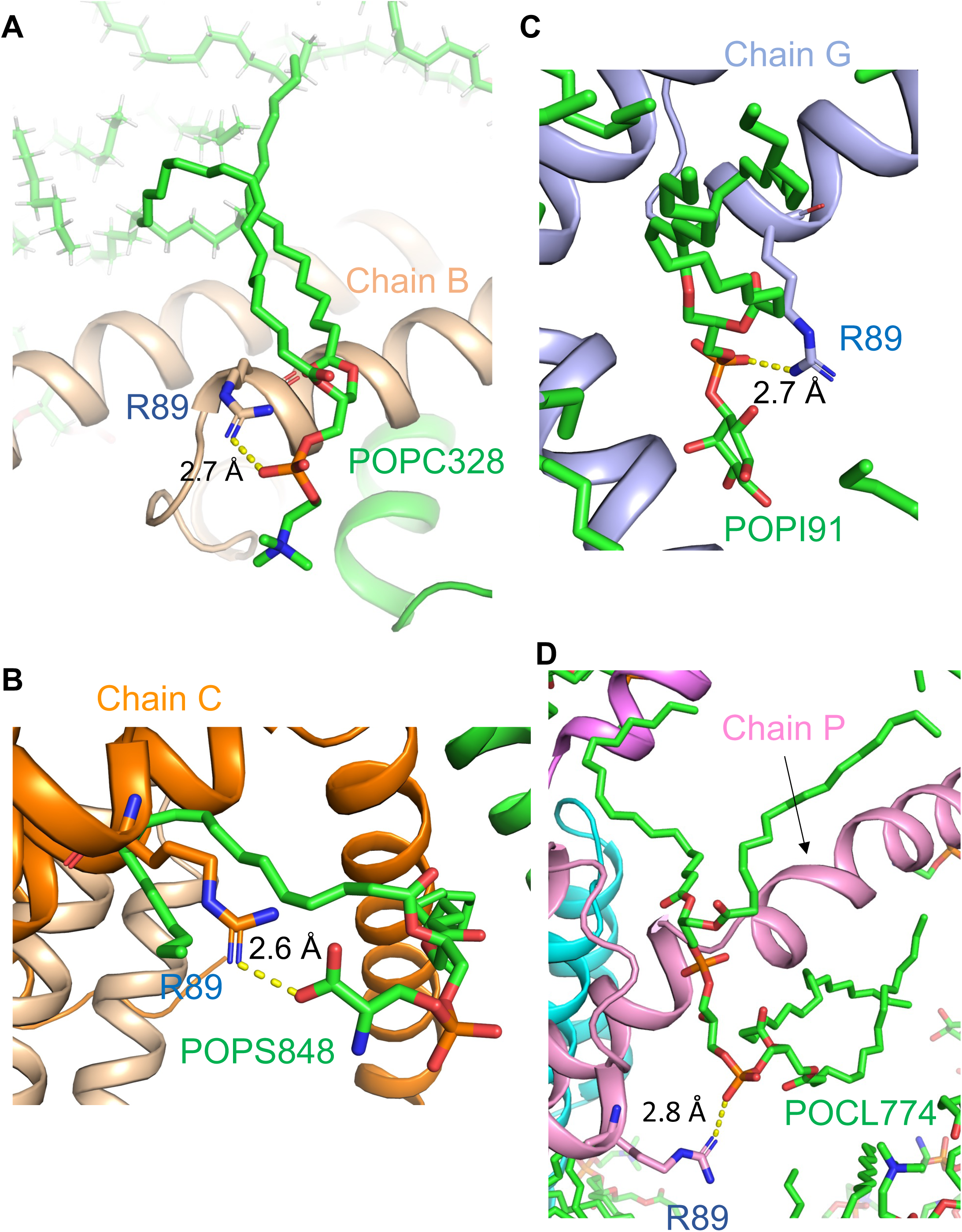
Lipids that interact with R89 in the pore wall observed by the end of MD simulations. Potential ionic or hydrogen bond between R89 and POPC (A), POPS (B), POPI (C) and POCL (D) is indicated by yellow dashed lines.

**Figure S11.**
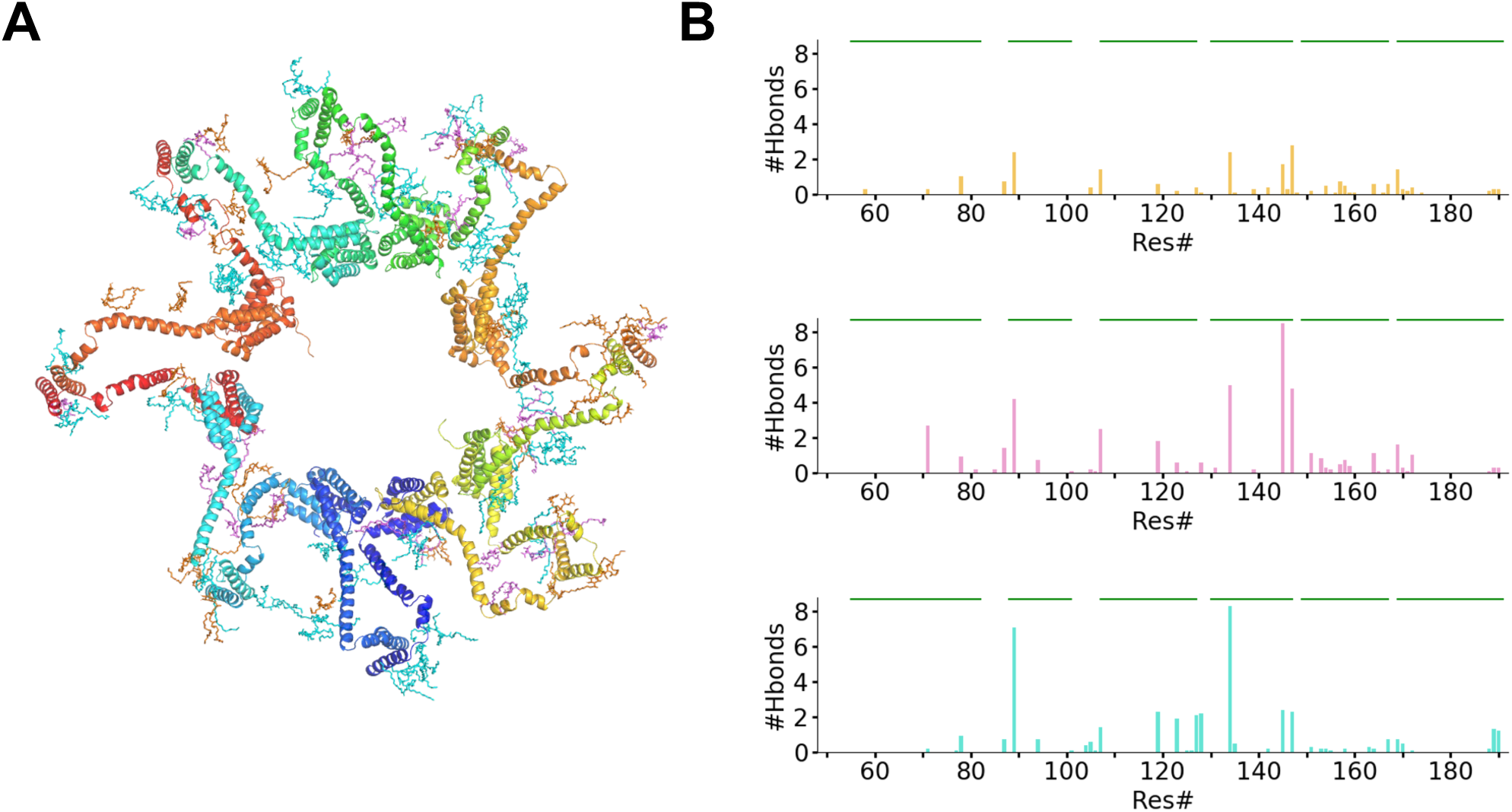
Protein-lipid interactions observed around a pore observed by the end of MD simulations. **A.** A snapshot of negatively charge lipids (PI in orange, PS in pink, and CL in cyan) within 10 Å of C_α_ of protein residues. **B.** Major H-bond between Bax residues and negatively charged lipids (PI in top, PS in middle, and CL in bottom graphs).

**Figure S12.**
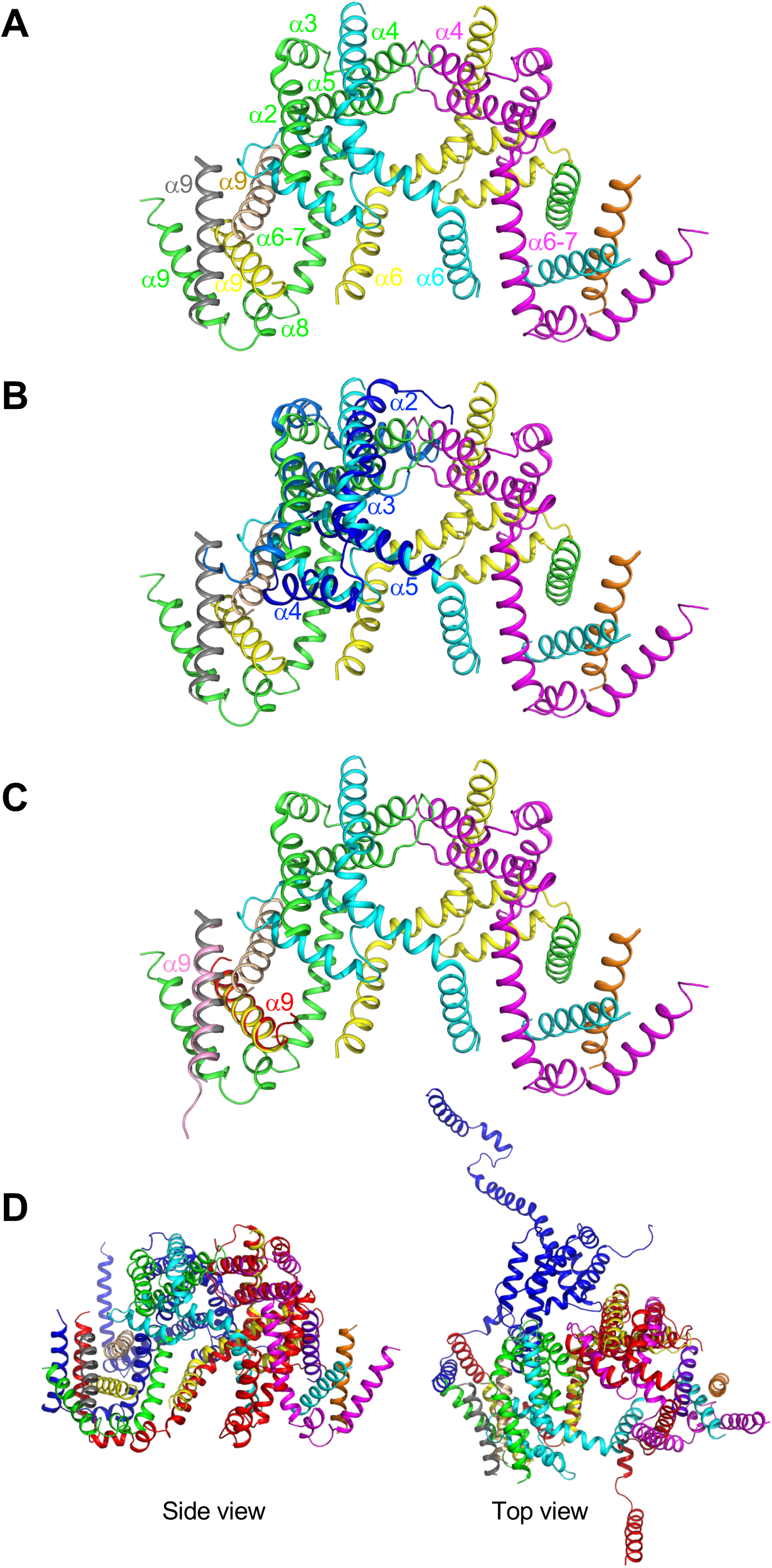
Comparison of Bax oligomer structure from cryoEM with α2-5 dimer and α9 dimer structures from NMR and Bax pore structure from MD simulations. **A.** CryoEM structure of a repeating unit for Bax oligomer (PDB ID: 9IXU) with protomers in different colors. The α-helices in green protomer are labeled. Of the green protomer, the α2-5 regions interact with the same regions in cyan protomer forming a BH3-in-groove dimer core. The α6 and α7 are merged into a single helix. The α9 and other three α9 helices, one from yellow protomer in this repeating unit, and two (gray and wheat) from an adjacent repeating unit, form a four-helical bundle. Each repeating unit contains two asymmetric dimers. The inter-dimer contacts are made by the α4 helices from green and magenta protomers, and by the α6 helices from green and yellow protomers and from magenta and cyan protomers. **B.** Superimposing the NMR structure (PDB ID: 6L8V) of α2-5 dimer in blue and marine with the cryoEM structure of α2-5 dimer in a repeating unit in green and cyan. The structure of the other α2-5 dimer in the repeating unit can also be superimposed by the NMR structure. **C.** Superimposing the NMR structure (PDBID: 6L95) of α9 dimer in red and pink with the the cryoEM structure of gray and yellow α9 helices in the four-α9 bundle on one end of the repeating unit. The structure of cyan and orange α9 helices on the other end of the repeating unit can also be superimposed by the NMR structure. **D.** Superimposing the structure of two neighboring α2-9 dimers (in red and blue) from a Bax pore structure generated by MD simulations with the cryoEM structure of the repeating unit. Only the red α2-5 dimer in the MD simulated structure is partially aligned with the yellow and magenta α2-5 dimer in the cryoEM structure.

**Supplementary Table S1.** Hydrophobicity scales for the α9 residues involved in mutations.

| Residue | Kyte-Doolittle<sup>a</sup> | Wimley-White<sup>b</sup> | Hessa-Heijne<sup>c</sup> | Moon-Fleming<sup>d</sup> | Zhao-London<sup>e</sup> |
| --- | --- | --- | --- | --- | --- |
| Ile | 4.5 | 0.31 | -0.60 | -1.56 | 1.97 |
| Val | 4.2 | -0.07 | -0.31 | -0.78 | 1.46 |
| Ala | 1.8 | -0.17 | 0.11 | 0.00 | 0.38 |
| Gly | -0.4 | -0.01 | 0.74 | 1.72 | -0.19 |
| Thr | -0.7 | -0.14 | 0.52 | 1.78 | -0.32 |
| Tyr | -1.3 | 0.94 | 0.68 | -1.09 | 0.49 |

a. Kyte, J. & Doolittle, R. F. A simple method for displaying the hydropathic character of a protein. *J Mol Biol* 157, 105-132, doi:10.1016/0022-2836(82)90515-0 (1982).
b. Wimley, W. C. & White, S. H. Experimentally determined hydrophobicity scale for proteins at membrane interfaces. *Nat Struct Biol* 3, 842-848, doi:10.1038/nsb1096-842 (1996).
c. Hessa, T. et al. Recognition of transmembrane helices by the endoplasmic reticulum translocon. *Nature* 433, 377-381, doi:10.1038/nature03216 (2005).
d. Moon, C. P. & Fleming, K. G. Side-chain hydrophobicity scale derived from transmembrane protein folding into lipid bilayers. *Proc Natl Acad Sci U S A* 108, 10174-10177, doi:10.1073/pnas.1103979108 (2011).
e. Zhao, G. & London, E. An amino acid "transmembrane tendency" scale that approaches the theoretical limit to accuracy for prediction of transmembrane helices: relationship to biological hydrophobicity. *Protein Sci* 15, 1987-2001, doi:10.1110/ps.062286306 (2006).

**Supplementary Table S2.** Change of hydrophobicity scales due to the α9 mutations.

| Mutation | Kyte-Doolittle | Wimley-White | Hessa-Heijne | Moon-Fleming | Zhao-London |
| --- | --- | --- | --- | --- | --- |
| I175A | ↓2.7 | ↓0.48 | ↓0.71 | ↓1.56 | ↓1.59 |
| G179Y | ↓0.9 | ↑0.95 | ↑0.06 | ↑2.81 | ↑0.68 |
| I187A | ↓2.7 | ↓0.48 | ↓0.71 | ↓1.56 | ↓1.59 |
| T174A | ↑2.5 | ↓0.03 | ↑0.41 | ↑1.78 | ↑0.70 |
| T182I | ↑5.2 | ↑0.45 | ↑1.12 | ↑3.34 | ↑2.29 |
| T186V | ↑4.9 | ↑0.07 | ↑0.83 | ↑2.56 | ↑1.78 |

↑ indicates an increase of hydrophobic scale.
↓ indicates a decrease of hydrophobic scale.
Number indicates a relative change of hydrophobic scale
= |Hydrophobicity scale of wild-type residue – Hydrophobicity scale of mutated residue|

**Supplementary Table S3.** Inter-dimeric protein-protein contacts: residue pairs within 6 Å and the frequency of their occurrence during MD simulations.

| Contact pairs |  | Frequency |
| --- | --- | --- |
| P130 | T135 | 8 |
| S72 | C126 | 4 |
| C126 | L132 | 4 |
| C126 | E131 | 4 |
| R134 | W139 | 4 |
| T127 | T135 | 3 |
| C126 | K128 | 3 |
| R134 | T135 | 3 |
| P130 | W139 | 3 |
| S72 | M74 | 2 |
| P130 | I136 | 2 |
| P130 | E131 | 2 |
| P130 | L132 | 2 |
| M74 | M74 | 2 |
| M74 | Q77 | 2 |
| R134 | F143 | 2 |
| F165 | P168 | 2 |
| E69 | C126 | 1 |
| S72 | Q77 | 1 |
| D68 | A81 | 1 |
| M137 | W139 | 1 |
| Q77 | R78 | 1 |
| M74 | A81 | 1 |
| C126 | T135 | 1 |
| S72 | S72 | 1 |
| D68 | M74 | 1 |
| I133 | T135 | 1 |
| S72 | L122 | 1 |
| Q77 | C126 | 1 |
| E131 | W139 | 1 |

**Supplementary Table S4.** Stable hydrogen bonds (H-bonds) between Bax protein and negatively charged lipids, phosphatidylinositol (PI), phosphatidylserine (PS) and cardiolipin (CL). The higher frequency the more frequently the hydrogen bond is formed.

| Bax protein-PI H-bond |  | Bax protein-PS H-bond |  | Bax protein-CL H-bond |  |
| --- | --- | --- | --- | --- | --- |
| residue | frequency | residue | frequency | residue | frequency |
| K58 | 3 | D71 | 27 | D71 | 2 |
| D71 | 3 | R78 | 9 | Q77 | 1 |
| R78 | 10 | A81 | 2 | R78 | 9 |
| S87 | 7 | T85 | 2 | S87 | 7 |
| R89<sup>a</sup> | 24 | S87 | 14 | R89 | 71 |
| F105 | 4 | R89 | 42 | R94 | 7 |
| W107 | 14 | R94 | 7 | S101 | 1 |
| K119 | 6 | S101 | 1 | N104 | 4 |
| K123 | 2 | F105 | 2 | F105 | 6 |
| T127 | 4 | N106 | 1 | N106 | 1 |
| K128 | 1 | W107 | 25 | W107 | 14 |
| R134 | 24 | K119 | 18 | K119 | 23 |
| T135 | 1 | K123 | 6 | K123 | 19 |
| W139 | 3 | L125 | 1 | L125 | 1 |
| D142 | 4 | K128 | 6 | C126 | 1 |
| R145 | 17 | E131 | 3 | T127 | 21 |
| E146 | 3 | R134 | 50 | K128 | 22 |
| R147 | 28 | W139 | 2 | R134 | 83 |
| L148 | 1 | R145 | 85 | T135 | 5 |
| W151 | 2 | R147 | 48 | D142 | 2 |
| D154 | 5 | W151 | 11 | R145 | 24 |
| G156 | 1 | Q153 | 8 | R147 | 23 |
| G157 | 7 | D154 | 3 | W151 | 3 |
| W158 | 5 | Q155 | 2 | Q153 | 2 |
| D159 | 1 | G157 | 5 | D154 | 2 |
| G160 | 1 | W158 | 7 | Q155 | 1 |
| Y164 | 6 | D159 | 4 | W158 | 2 |
| G166 | 1 | Y164 | 11 | S163 | 3 |
| T167 | 6 | F165 | 1 | Y164 | 2 |
| T169 | 14 | T167 | 2 | T167 | 7 |
| W170 | 3 | T169 | 16 | T169 | 7 |
| Q171 | 2 | W170 | 3 | W170 | 5 |
| T172 | 4 | Q171 | 1 | T172 | 1 |
| T174 | 1 | T172 | 10 | W188 | 2 |
| W188 | 2 | W188 | 1 | K189 | 13 |
| K189 | 3 | K189 | 3 | K190 | 12 |
| K190 | 3 | K190 | 3 |  |  |

## References

1. Andrews DW (2014) Pores of no return. Mol Cell 56: 465–466

2. Andrews DW (2022) The case for brakes: Why restrain the size of Bax and Bak pores in outer mitochondrial membranes? Mol Cell 82: 882–883

3. Barnes CA, Mishra P, Baber JL, Strub MP, Tjandra N (2017) Conformational Heterogeneity in the Activation Mechanism of Bax. Structure 25: 1310–1316 e1313

4. Bartels C, Xia TH, Billeter M, Guntert P, Wuthrich K (1995) The program XEASY for computer-supported NMR spectral analysis of biological macromolecules. J Biomol NMR 6: 1–10

5. Best RB, Zhu X, Shim J, Lopes PE, Mittal J, Feig M, Mackerell AD, Jr. (2012) Optimization of the additive CHARMM all-atom protein force field targeting improved sampling of the backbone phi, psi and side-chain chi(1) and chi(2) dihedral angles. J Chem Theory Comput 8: 3257–3273

6. Bleicken S, Assafa TE, Stegmueller C, Wittig A, Garcia-Saez AJ, Bordignon E (2018) Topology of active, membrane-embedded Bax in the context of a toroidal pore. Cell Death Differ 25: 1717–1731

7. Bleicken S, Jeschke G, Stegmueller C, Salvador-Gallego R, Garcia-Saez AJ, Bordignon E (2014) Structural model of active Bax at the membrane. Mol Cell 56: 496–505

8. Brielle ES, Arkin IT (2020) Quantitative Analysis of Multiplex H-Bonds. J Am Chem Soc 142: 14150–14157

9. Chi X, Nguyen D, Pemberton JM, Osterlund EJ, Liu Q, Brahmbhatt H, Zhang Z, Lin J, Leber B, Andrews DW (2020) The carboxyl-terminal sequence of bim enables bax activation and killing of unprimed cells. Elife 9

10. Chio US, Cho H, Shan SO (2017) Mechanisms of Tail-Anchored Membrane Protein Targeting and Insertion. Annu Rev Cell Dev Biol 33: 417–438

11. Cosentino K, Garcia-Saez AJ (2017) Bax and Bak Pores: Are We Closing the Circle? Trends Cell Biol 27: 266–275

12. Cosentino K, Hertlein V, Jenner A, Dellmann T, Gojkovic M, Pena-Blanco A, Dadsena S, Wajngarten N, Danial JSH, Thevathasan JV et al (2022) The interplay between BAX and BAK tunes apoptotic pore growth to control mitochondrial-DNA-mediated inflammation. Mol Cell 82: 933–949 e939

13. Curran AR, Engelman DM (2003) Sequence motifs, polar interactions and conformational changes in helical membrane proteins. Curr Opin Struct Biol 13: 412–417

14. Czabotar PE, Westphal D, Dewson G, Ma S, Hockings C, Fairlie WD, Lee EF, Yao S, Robin AY, Smith BJ et al (2013) Bax crystal structures reveal how BH3 domains activate Bax and nucleate its oligomerization to induce apoptosis. Cell 152: 519–531

15. D.A. Case IYB-S, S.R. Brozell, D.S. Cerutti, T.E. Cheatham, III, V.W.D. Cruzeiro, T.A. Darden, R.E. Duke DG, M.K. Gilson, H. Gohlke, A.W. Goetz, D. Greene, R Harris, N. Homeyer, Y. Huang,, S. Izadi AK, T. Kurtzman, T.S. Lee, S. LeGrand, P. Li, C. Lin, J. Liu, T. Luchko, R. Luo, D.J., Mermelstein KMM, Y. Miao, G. Monard, C. Nguyen, H. Nguyen, I. Omelyan, A. Onufriev, F. Pan, R., Qi DRR, A. Roitberg, C. Sagui, S. Schott-Verdugo, J. Shen, C.L. Simmerling, J. Smith, R. Salomon-, Ferrer JS, R.C. Walker, J. Wang, H. Wei, R.M. Wolf, X. Wu, L. Xiao, D.M. York and P.A. Kollman (2018) AMBER 2018, University of California, San Francisco.

16. Dal Peraro M, van der Goot FG (2016) Pore-forming toxins: ancient, but never really out of fashion. Nat Rev Microbiol 14: 77–92

17. Delaglio F, Grzesiek S, Vuister GW, Zhu G, Pfeifer J, Bax A (1995) NMRPipe: a multidimensional spectral processing system based on UNIX pipes. J Biomol NMR 6: 277–293

18. Dengler MA, Robin AY, Gibson L, Li MX, Sandow JJ, Iyer S, Webb AI, Westphal D, Dewson G, Adams JM (2019) BAX Activation: Mutations Near Its Proposed Non-canonical BH3 Binding Site Reveal Allosteric Changes Controlling Mitochondrial Association. Cell Rep 27: 359–373 e356

19. Dewson G, Ma S, Frederick P, Hockings C, Tan I, Kratina T, Kluck RM (2012) Bax dimerizes via a symmetric BH3:groove interface during apoptosis. Cell Death Differ 19: 661–670

20. Ding J, Mooers BHM, Zhang Z, Kale J, Falcone D, McNichol J, Huang B, Zhang XC, Xing C, Andrews DW et al (2014) After embedding in membranes antiapoptotic Bcl-XL protein binds both Bcl-2 homology region 3 and helix 1 of proapoptotic Bax protein to inhibit apoptotic mitochondrial permeabilization. J Biol Chem 289: 11873–11896

21. Ding J, Zhang Z, Roberts GJ, Falcone M, Miao Y, Shao Y, Zhang XC, Andrews DW, Lin J (2010) Bcl-2 and Bax interact via the BH1-3 groove-BH3 motif interface and a novel interface involving the BH4 motif. J Biol Chem 285: 28749–28763

22. Dlugosz PJ, Billen LP, Annis MG, Zhu W, Zhang Z, Lin J, Leber B, Andrews DW (2006) Bcl-2 changes conformation to inhibit Bax oligomerization. EMBO J 25: 2287–2296

23. Edlich F, Banerjee S, Suzuki M, Cleland MM, Arnoult D, Wang C, Neutzner A, Tjandra N, Youle RJ (2011) Bcl-x(L) retrotranslocates Bax from the mitochondria into the cytosol. Cell 145: 104–116

24. Escobedo A, Topal B, Kunze MBA, Aranda J, Chiesa G, Mungianu D, Bernardo-Seisdedos G, Eftekharzadeh B, Gairi M, Pierattelli R et al (2019) Side chain to main chain hydrogen bonds stabilize a polyglutamine helix in a transcription factor. Nat Commun 10: 2034

25. Eswar N, Ramakrishnan C (2000) Deterministic features of side-chain main-chain hydrogen bonds in globular protein structures. Protein Eng 13: 227–238

26. Feldblum ES, Arkin IT (2014) Strength of a bifurcated H bond. Proc Natl Acad Sci U S A 111: 4085–4090

27. Fink A, Sal-Man N, Gerber D, Shai Y (2012) Transmembrane domains interactions within the membrane milieu: principles, advances and challenges. Biochim Biophys Acta 1818: 974–983

28. Gahl RF, He Y, Yu S, Tjandra N (2014) Conformational rearrangements in the pro-apoptotic protein, Bax, as it inserts into mitochondria: a cellular death switch. J Biol Chem 289: 32871–32882

29. Gao J, Bosco DA, Powers ET, Kelly JW (2009) Localized thermodynamic coupling between hydrogen bonding and microenvironment polarity substantially stabilizes proteins. Nat Struct Mol Biol 16: 684–690

30. Garg P, Nemec KN, Khaled AR, Tatulian SA (2013) Transmembrane pore formation by the carboxyl terminus of Bax protein. Biochim Biophys Acta 1828: 732–742

31. Garner TP, Reyna DE, Priyadarshi A, Chen HC, Li S, Wu Y, Ganesan YT, Malashkevich VN, Cheng EH, Gavathiotis E (2016) An Autoinhibited Dimeric Form of BAX Regulates the BAX Activation Pathway. Mol Cell 63: 485–497

32. Gavathiotis E, Reyna DE, Davis ML, Bird GH, Walensky LD (2010) BH3-triggered structural reorganization drives the activation of proapoptotic BAX. Mol Cell 40: 481–492

33. Gavathiotis E, Suzuki M, Davis ML, Pitter K, Bird GH, Katz SG, Tu HC, Kim H, Cheng EH, Tjandra N et al (2008) BAX activation is initiated at a novel interaction site. Nature 455: 1076–1081

34. Hessa T, Kim H, Bihlmaier K, Lundin C, Boekel J, Andersson H, Nilsson I, White SH, von Heijne G (2005) Recognition of transmembrane helices by the endoplasmic reticulum translocon. Nature 433: 377–381

35. Hunter JD (2007) Matplotlib: A 2D Graphics Environment. Computing in Science & Engineering 9: 90–95

36. Iyer S, Uren RT, Dengler MA, Shi MX, Uno E, Adams JM, Dewson G, Kluck RM (2020) Robust autoactivation for apoptosis by BAK but not BAX highlights BAK as an important therapeutic target. Cell Death Dis 11: 268

37. Jo S, Kim T, Iyer VG, Im W (2008) CHARMM-GUI: a web-based graphical user interface for CHARMM. J Comput Chem 29: 1859–1865

38. Kale J, Kutuk O, Brito GC, Andrews TS, Leber B, Letai A, Andrews DW (2018a) Phosphorylation switches Bax from promoting to inhibiting apoptosis thereby increasing drug resistance. EMBO Rep 19

39. Kale J, Osterlund EJ, Andrews DW (2018b) BCL-2 family proteins: changing partners in the dance towards death. Cell Death Differ 25: 65–80

40. Kuwana T, King LE, Cosentino K, Suess J, Garcia-Saez AJ, Gilmore AP, Newmeyer DD (2020) Mitochondrial residence of the apoptosis inducer BAX is more important than BAX oligomerization in promoting membrane permeabilization. J Biol Chem 295: 1623–1636

41. Kuwana T, Olson NH, Kiosses WB, Peters B, Newmeyer DD (2016) Pro-apoptotic Bax molecules densely populate the edges of membrane pores. Sci Rep 6: 27299

42. Kyrychenko A, Vasquez-Montes V, Ladokhin AS (2022) Advantages of Quantitative Analysis of Depth-Dependent Fluorescence Quenching: Case Study of BAX. J Membr Biol 255: 461–468

43. Kyte J, Doolittle RF (1982) A simple method for displaying the hydropathic character of a protein. J Mol Biol 157: 105–132

44. Liao C, Zhang Z, Kale J, Andrews DW, Lin J, Li J (2016) Conformational Heterogeneity of Bax Helix 9 Dimer for Apoptotic Pore Formation. Sci Rep 6: 29502

45. Lovell JF, Billen LP, Bindner S, Shamas-Din A, Fradin C, Leber B, Andrews DW (2008) Membrane binding by tBid initiates an ordered series of events culminating in membrane permeabilization by Bax. Cell 135: 1074–1084

46. Luna-Vargas MPA, Chipuk JE (2016) Physiological and Pharmacological Control of BAK, BAX, and Beyond. Trends Cell Biol 26: 906–917

47. Lv F, Qi F, Zhang Z, Wen M, Kale J, Piai A, Du L, Wang S, Zhou L, Yang Y et al (2021) An amphipathic Bax core dimer forms part of the apoptotic pore wall in the mitochondrial membrane. EMBO J 40: e106438

48. Moldoveanu T, Czabotar PE (2020) BAX, BAK, and BOK: A Coming of Age for the BCL-2 Family Effector Proteins. Cold Spring Harb Perspect Biol 12

49. Moon CP, Fleming KG (2011) Side-chain hydrophobicity scale derived from transmembrane protein folding into lipid bilayers. Proc Natl Acad Sci U S A 108: 10174–10177

50. Roe DR, Cheatham TE, 3rd (2013) PTRAJ and CPPTRAJ: Software for Processing and Analysis of Molecular Dynamics Trajectory Data. J Chem Theory Comput 9: 3084–3095

51. Ruffolo SC, Shore GC (2003) BCL-2 selectively interacts with the BID-induced open conformer of BAK, inhibiting BAK auto-oligomerization. J Biol Chem 278: 25039–25045

52. Salzmann M, Wider G, Pervushin K, Senn H, Wüthrich K (1999) TROSY-type Triple-Resonance Experiments for Sequential NMR Assignments of Large Proteins. Journal of the American Chemical Society 121: 844–848

53. Scharnagl C, Pester O, Hornburg P, Hornburg D, Gotz A, Langosch D (2014) Side-chain to main-chain hydrogen bonding controls the intrinsic backbone dynamics of the amyloid precursor protein transmembrane helix. Biophys J 106: 1318–1326

54. Spitz AZ, Gavathiotis E (2022) Physiological and pharmacological modulation of BAX. Trends Pharmacol Sci 43: 206–220

55. Sung TC, Li CY, Lai YC, Hung CL, Shih O, Yeh YQ, Jeng US, Chiang YW (2015) Solution Structure of Apoptotic BAX Oligomer: Oligomerization Likely Precedes Membrane Insertion. Structure 23: 1878–1888

56. Suzuki M, Youle RJ, Tjandra N (2000) Structure of Bax: coregulation of dimer formation and intracellular localization. Cell 103: 645–654

57. Tan C, Dlugosz PJ, Peng J, Zhang Z, Lapolla SM, Plafker SM, Andrews DW, Lin J (2006) Auto-activation of the apoptosis protein Bax increases mitochondrial membrane permeability and is inhibited by Bcl-2. J Biol Chem 281: 14764–14775

58. Teese MG, Langosch D (2015) Role of GxxxG Motifs in Transmembrane Domain Interactions. Biochemistry 54: 5125–5135

59. Tsai CJ, Liu S, Hung CL, Jhong SR, Sung TC, Chiang YW (2015) BAX-induced apoptosis can be initiated through a conformational selection mechanism. Structure 23: 139–148

60. Vogel S, Raulf N, Bregenhorn S, Biniossek ML, Maurer U, Czabotar P, Borner C (2012) Cytosolic Bax: does it require binding proteins to keep its pro-apoptotic activity in check? J Biol Chem 287: 9112–9127

61. Vranken WF, Boucher W, Stevens TJ, Fogh RH, Pajon A, Llinas M, Ulrich EL, Markley JL, Ionides J, Laue ED (2005) The CCPN data model for NMR spectroscopy: development of a software pipeline. Proteins 59: 687–696

62. Walensky LD (2019) Targeting BAX to drug death directly. Nat Chem Biol 15: 657–665

63. Westphal D, Dewson G, Menard M, Frederick P, Iyer S, Bartolo R, Gibson L, Czabotar PE, Smith BJ, Adams JM et al (2014) Apoptotic pore formation is associated with in-plane insertion of Bak or Bax central helices into the mitochondrial outer membrane. Proc Natl Acad Sci U S A 111: E4076–4085

64. Wimley WC, White SH (1996) Experimentally determined hydrophobicity scale for proteins at membrane interfaces. Nat Struct Biol 3: 842–848

65. Zhang Y, Tian L, Huang G, Ge X, Kong F, Wang P, Xu Y, Shi Y (2025) Structural basis of BAX pore formation. Science 388: eadv4314

66. Zhang Z, Subramaniam S, Kale J, Liao C, Huang B, Brahmbhatt H, Condon SG, Lapolla SM, Hays FA, Ding J et al (2016) BH3-in-groove dimerization initiates and helix 9 dimerization expands Bax pore assembly in membranes. EMBO J 35: 208–236

67. Zhang Z, Zhu W, Lapolla SM, Miao Y, Shao Y, Falcone M, Boreham D, McFarlane N, Ding J, Johnson AE et al (2010) Bax forms an oligomer via separate, yet interdependent, surfaces. J Biol Chem 285: 17614–17627

68. Zhao G, London E (2006) An amino acid "transmembrane tendency" scale that approaches the theoretical limit to accuracy for prediction of transmembrane helices: relationship to biological hydrophobicity. Protein Sci 15: 1987–2001

